# Elucidating the viral and host factors enabling the cross-species transmission of primate lentiviruses from simians to humans

**DOI:** 10.1101/2020.10.13.337303

**Authors:** Denis M. Tebit, Gabrielle Nickel, Richard Gibson, Crystal Carpenter, Myriam Rodriguez, Nicolas J. Hathaway, Katie Bain, Angel L. Reyes-Rodriguez, Jennifer Bonogo, David Canaday, David McDonald, Jeffrey A. Bailey, Eric J. Arts

## Abstract

The HIV-1 epidemic originated from a cross-species transmission of a primate lentivirus from chimpanzees to humans near the turn of the 18^th^ century. Simian immunodeficiency viruses have been jumping between old world monkeys in West/Central Africa for thousands of years. So why did HIV-1 only emerge in the past century? This study examined the replicative fitness, transmission, restriction, and cytopathogenicity of 26 primate lentiviruses. Pairwise competitions of these primate lentiviruses revealed that SIVcpz had the highest replicative fitness in human or chimpanzee peripheral blood mononuclear cells, even higher fitness than HIV-1 group M strains responsible for 37 million infections worldwide. In contrast the “HIV-2 lineage” (SIVsmm, SIVmac, SIVagm, and HIV-2) had the lowest replicative fitness. SIVcpz strains were less inhibited by human restriction factors than the “HIV-2 lineage” strains, a restriction that was inversely correlated with replicative fitness. SIVcpz from the chimpanzee subspecies *Pan troglodytes troglodytes* (*Ptt*) was slightly more fit in human cells than the strains from *Pt schweinfurthii* (*Pts*). However, unlike all other primate lentiviruses (including the HIV-2 lineage), SIVcpz was nonpathogenic in human tonsillar tissue and did not deplete CD4+ T-cells, consistent with the slow or nonpathogenic disease observed in chimpanzees. Despite the close phylogenetic relationship between SIVcpz_Ptt and HIV-1, this epidemic was either caused by cross species transmission of a rare, undiscovered SIVcpz strain of higher virulence or higher virulence differentially evolved among HIV-1 subtypes during the human epidemic.

**Author summary:** Invasion of wild animal habitats by humans can have devastating consequences for the human population as evident by the HIV-1 and SARS-CoV-2 epidemics. With SARS-CoV-2, a recent zoonotic jump, likely from bats, will help to identify a coronavirus progenitor. In contrast, simian immunodeficiency virus (SIV) jumped into humans over 100 years ago from a possibly extinct sub-species of chimpanzees and/or extinct lineage of SIV. We examined replicative fitness and pathogenesis of 26 different primate lentiviruses in human and chimpanzee primary lymphoid cells from blood and within tonsils. SIV from a specific chimpanzee species and lowland gorillas were the most capable of infecting and replicating in human and chimp lymphoid cells but they did not result in the pathogenesis related to disease in humans. In contrast, SIV from other old world monkeys were pathogenic but could not replicate efficiently in human cells. We propose the main HIV-1 is derived from a distinct jump of a very rare SIV strain in chimps leading to AIDS pandemic.

## Introduction

Simian immunodeficiency viruses (SIV) are a collection of primate lentiviruses differing as much as 50% (in nucleotide diversity) and infecting a genetically diverse population of old world monkeys(1,2). In contrast the distinction between “human” immunodeficiency virus type 1 (HIV-1) and “simian”IV from chimpanzees (cpz) are more in the assigned name considering SIVcpz and HIV-1 share more homology than SIVs that infected different old world monkeys(1–4). Zoonotic transmissions of SIV occurred through several independent jumps from chimpanzees to humans leading to the establishment of 4 groups within HIV-1 (M, N, O and P) and from sooty mangabeys to humans for the 9 groups (A-I) within HIV-2(1–7). HIV-1 group M is the most prevalent human lentivirus accounting for >95% of the 36 million infections in humans while HIV-2 and HIV-1 group O are responsible for 1-2 million and <30,000 infections, respectively(2,8). The rare HIV-1 groups N and P are found in less than a hundred individuals residing in or visiting Equatorial rainforest of West and Central Africa(2,9–13), also the origin of HIV-1 group M zoonotic jump. West Central African countries with Cameroon at its epicenter, represent the oldest epidemic illustrated by high HIV genetic variability in the current infected population(2,13).

There are more than 40 different SIV lineages infecting more than 45 species of African non-human primates, most commonly chimpanzees, gorillas and other old world monkeys(3–5,14–16). As an example, SIVsmm infects sooty mangabeys (smm) (or *Cercocebus atys)* in West Africa and is the likely source of HIV-2(17–19). SIVcpz infects the *Pan troglodytes troglodytes* (*Ptt*) and *Pan troglodytes schweinfurthii* (*Pts*) sub-species of chimpanzees common in central and west central Africa(7,20–22). While SIVcpz from *Ptt* is the origin of HIV-1 groups M and N, *Pts* related SIV lineages have not been found in humans(7,122,23). Likewise, SIVcpz has not jumped to other subspecies of chimpanzees (i.e. *Pt verus or vellerosus or* bonomos) which raises the possibility of incompatibility or lack of opportunity for this cross-species transmission(24–27). It is still unclear if SIV infecting lowland gorillas (*Gorilla gorilla or gor*) inhabiting the Equatorial forest of Central Africa(15,16) were the direct link for the primate lentiviral jump to humans to establish HIV-1 groups O and P or if this jump involved chimpanzees as an intermediate host.

A cross-species lentiviral inoculation establishing a new, successful zoonosis may be governed by higher replicative capacity(2,28) and reduced sensitivity to innate restriction factors in the new primate host(22,29). As an example, increased sensitivity to human restriction factors or low replicative fitness of SIVcpz from *Pts* chimpanzees in susceptible human CD4+ cells may have prevented this cross species transmission. Alternatively, humans were never exposed to an SIV from infected *Pts* chimpanzees. Given the deforestation and exploitation in sub-Saharan Africa after the turn of the 18^th^ century, a time which coincided with the introduction then rapid expansions of the HIV-1 and −2 epidemics, it is unlikely but not impossible that humans were only exposed to SIV from *Ptt* and not *Pts* chimpanzees. Primate lentiviruses have been jumping and circulating through old world primate populations, including humans, for thousands of years(30). Cross-specific transmission and variable host adaptions likely resulted in extinction of many primate lentiviral lineages, attenuations, and even the birth of novel lineages through recombination events that occur during super-infections with SIVs of different primates(31–34). In fact, SIVcpz are recombinants between two lentiviruses originating from the red capped mangabey and greater spot-nosed monkey or a closely related species(35).

Following the zoonotic transmission to humans around the 1920s, HIV-1 group M rapidly evolved into 10 different subtypes within the Congo Basin but with increased human migration and trade with sub-Saharan Africa, many HIV-1 subtypes founded new epidemics across the globe and even recombined to generate new circulating recombinant forms (CRFs)(2)(1,36). This collection of group M subtypes have evolved almost equidistant from the root of the HIV tree but despite the similar evolutionary rates, considerable differences in virulence and transmission efficiency may still be shaping subtype spread(37–39). In particular, HIV-1 subtype D may be the most virulent in humans(37,40,41) but has decreased in prevalence in the past 30 years whereas the rapid expansion and predominance of HIV-1 subtype C worldwide may be related to its lower virulence and average transmissibility(2). Following thousands of pairwise competitions of HIV-1 in human peripheral blood mononuclear cells (PBMCs), primary T cells, macrophages, tonsillar tissue, and DC-T cell cocultures, we have established a rank order in replicative or “pathogenic” fitness: HIV-1 groupM/subtype F ≥ B = D = CRF12,17_BF > A = CRF02_AG = CRF01_AE > C > HIV-1 grpO > HIV-2(38,42,43). Although HIV-1 replicative fitness in primary T cells and macrophages is a strong correlate of disease progression in humans(37,44,45), this relationship is not absolute especially with HIV-2 in humans and some SIV strains in African green monkeys (agm) and chimpanzees(46–52). A better surrogate of virulence may be direct infection of primary lymphoid organs like tonsillar tissue in which viral pathogenicity, T-cell depletion, and viral propagation rates can be measured(53,54).

In the present study, we utilized various primary human cell types to determine the *ex vivo* fitness and pathogenicity of a diverse set of 21 primate lentiviral isolates that included HIV-1 groups M, N, O, HIV-2, and SIVs of cpz, gor, mac, smm, and agm. We have performed over 300 pairwise dual infections using the 21 primate lentivirus isolates in peripheral blood mononuclear cells from both human and chimp (*Pan troglodytes verus; Ptv*) donors, along with monoinfection of PMBCs and cell lines, and finally dendritic cell-mediated transinfections of human (Homo sapiens; *Hs*) PBMCs. We determine the relative repression of primate lentivirus replication in human cells by various restriction factors and correlated this to replicative fitness. Finally, we accessed the cytopathogenicity on CD4+ T cells in human tonsillar explant tissue following infection with the HIV-1, HIV-2 and SIV isolates. Our results show that SIV strains from chimps and gorillas had similar *ex vivo* fitness as HIV-1 group M strains in both human and chimp PBMCs, while HIV-1 group O, SIVmac/smm, and HIV-2 isolates had sequentially lower fitness. Based on the higher relative fitness and reduced restriction on SIVcpz-*Ptt* in human T-cells compared to that of SIVcpz-*Pts* or any other SIV strain, we provide experimental evidence that SIVcpz from *Ptt* was most adapt in establishing infection in humans. However, this initial zoonotic transmission to humans was likely non-pathogenic based on tonsillar tissue studies.

## RESULTS

### HIV-1, HIV-2 and SIV replication in human PBMCs and U87 cell line

Twenty-six HIV and SIV strains representing 10 HIV-1 group M, 3 group O, 1 group N, 6 SIVcpz, 1 SIVgor, 2 HIV-2 and 1 each of SIVsmm, SIVmac and SIVagm (Figure 1A and B) were propagated on either U87.CD4.CCR5 cell line or human PBMC. Infectious titer was determined in human PBMCs using a standard TCID_50_ assay (Figure 1B). All isolates were R5 as confirmed by their replication on U87.CD4.CCR5 but not U87CXCR4 cell lines. Genotypes were confirmed through Sanger sequencing and phylogenetic analyses of gag and env; neighbor-joining trees shown in Figure 1A and Supplementary Figure 1. The group M strains clustered more closely with SIVcpz MB897 (Fig 1A) while SIVgor_CP684 grouped with the group O strains, as previously described (Fig 1A and B). SIVcpz EK505 is more closely related to the HIV-1 group N strain, YBF30. The HIV-2 strains VI1835 and VI1905 (both subtype A) clustered with SIVsmm, the source of HIV-2 (Fig 1A; Supplementary Figure 1).

**Figure 1:**
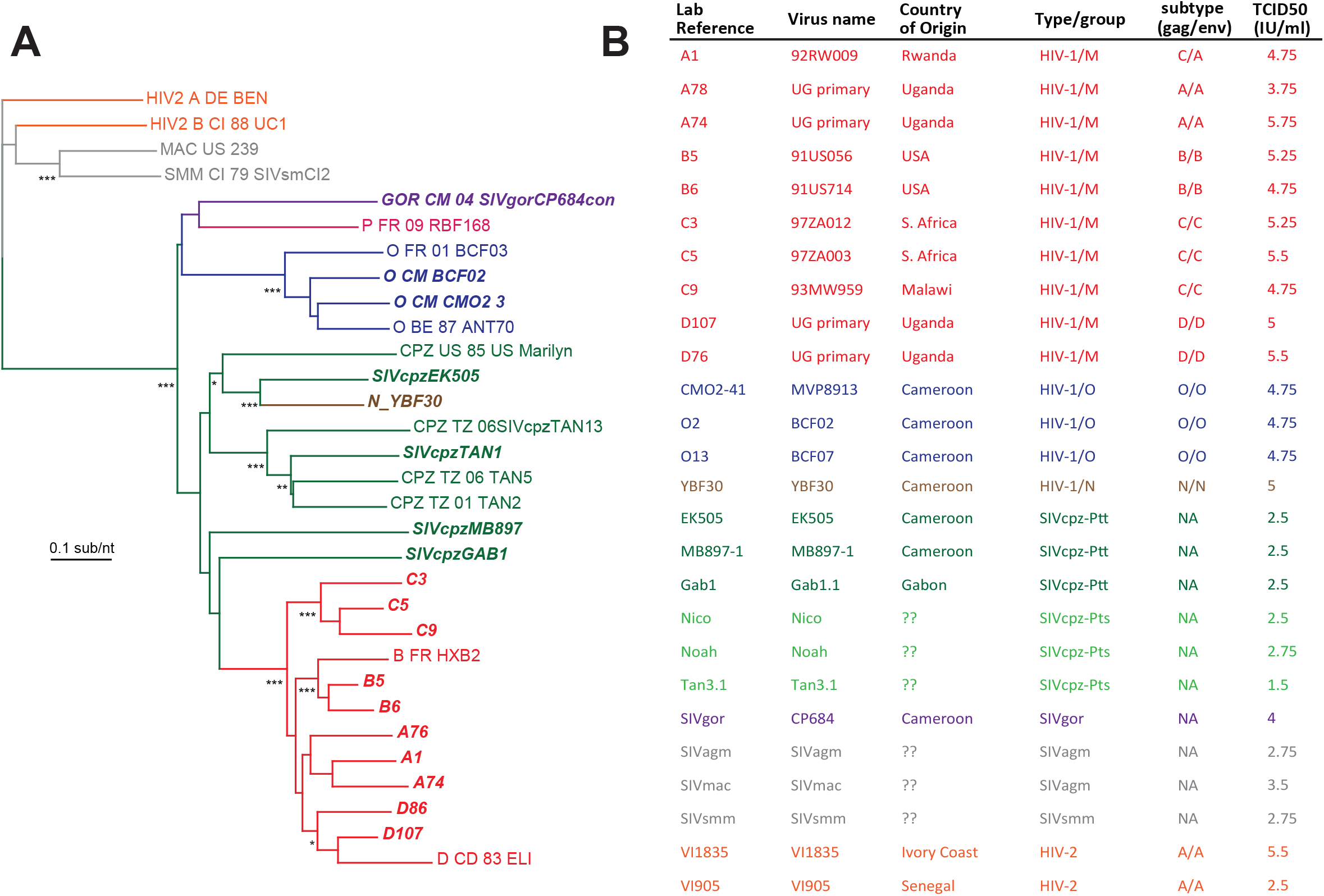
Characteristics of lentiviruses used in this study. (A) Neighbor joining phylogenetic tree of the *env* sequence of primate lentiviruses with reference strains. (B) List of study viruses showing origin, genotype, and virus titres (tissue culture infectious dose for 50% infectivity; TCID_50_).

The replication kinetics of all the primate lentiviruses were determined on human PBMCs and the brain glioma cell line U87 expressing huCD4 and either from human (*Hs* for *Homo sapiens*) CCR5 or rhesus macaque (*Mm* for *Macaca mulatta*) CCR5. All cell types were infected with the same MOI and monitored for virus production (and cell viability, see below) for 12 days to determine the amount of reverse transcriptase (Methods). Virus replication kinetics over a course of a 12 day infection in PHA/IL2-treated human PBMCs (as measured by area under the RT activity curve) were not significantly different between the HIV-1 group M, O and SIVcpz strains (Fig 2A). In the human brain glioma cell line expressing *Hs*CD4 with *Hs*CCR5 or *Mm*CCR5, the HIV-1 group M strains had 10-fold and nearly 100-fold greater replication kinetics than HIV-1 group O and SIVcpz strains, respectively (Figure 2A). When comparing growth kinetics for specific primate lentivirus strains, the HIV-1 group M subtype A strain, A74 (Fig 2Bi) replicates more efficiently in the U87 cell lines than in human PBMCs whereas the human lentiviruses, HIV-1 group O CM02-41 (Fig 2Bii) and HIV-2 VI1835 isolates (Fig 2Biii) replicate to similar levels in PBMCs and the U87 cell lines (with *Hs* or *Mm*CCR5). However, the SIV strains of chimpanzees (cpz) (Supplemental figure 2) and of African green monkeys (agm) (Fig 2Biv) replicated at higher levels in *Hs*PBMCs than in U87 cells. We propose that the HIV-1 group M is more adapt to replication in many human cell types aside from just CD4+ T-lymphocytes as compared to HIV-1 group O and HIV-2.

**Figure 2:**
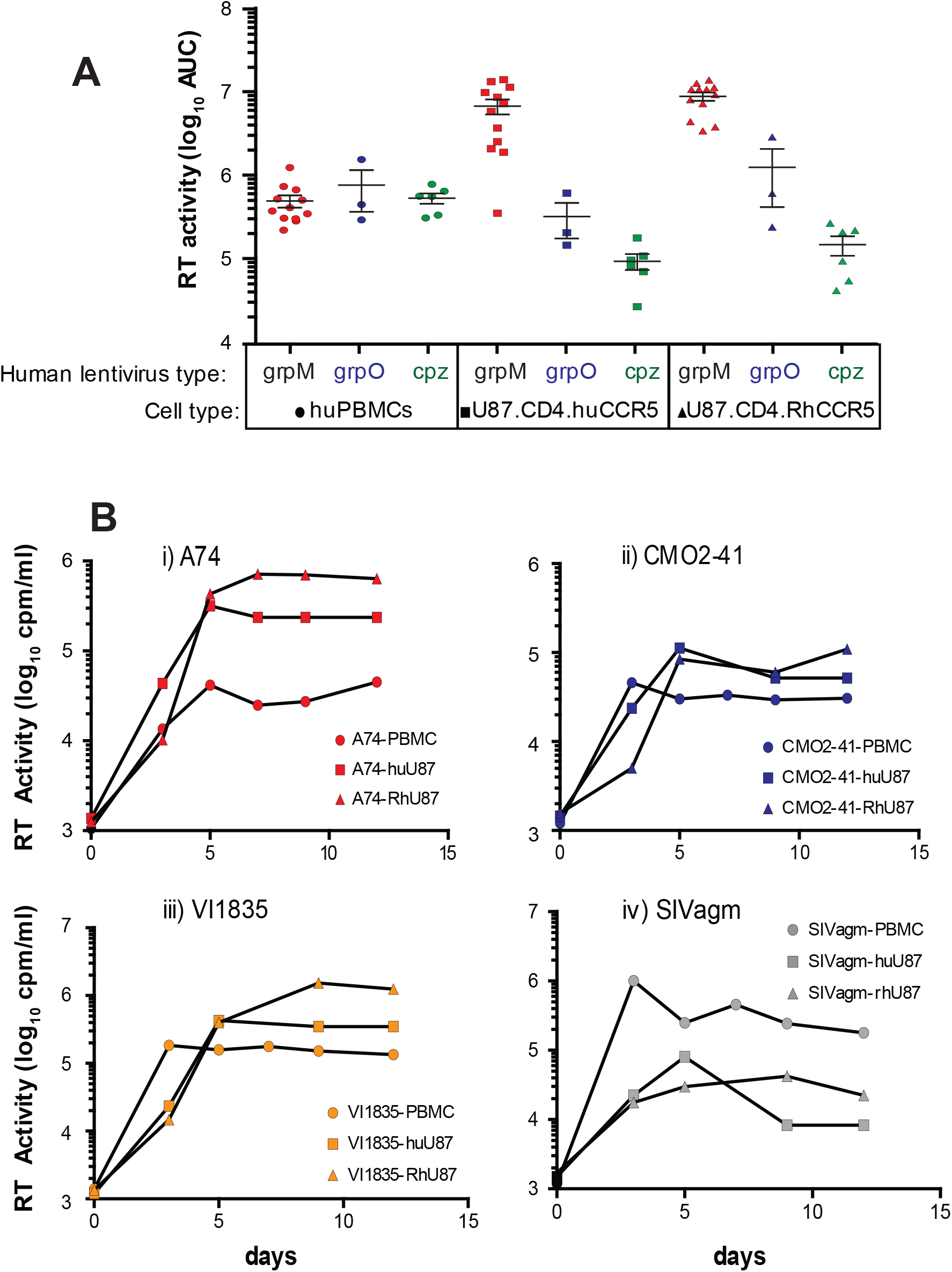
Replication of primate lentiviruses in human PBMCs or CD4+ U87 cells expressing human or rhesus macaque CCR5 as measured by reverse transcriptase activity in supernatant. Panel (A) shows area-under-the curve of virus production for HIV-1 group M, group O, and SIVcpz strains over a 12 day infection as measured by RT activity. Examples of individual production of group M A74 (i), group O CMO2-41 (ii), HIV-2 VI1835 (iii), and SIVagm (iv) is shown. All infections were done in triplicate. Standard deviations at each time point were within 20% of the value on average and are not shown.

### Dendritic cells enhance HIV and SIV infection in PBMCs

Within a human host, activated CD4+ T-cells in the gut, lymph nodes, and other tissues may be responsible for the majority of HIV-1 replication. T-cell activation and delivery of the HIV may be mediated by direct dendritic cell contact. The replication of primate lentiviruses in PBMCs mediated by human MDDC trans-infection was significantly greater (2- to 3-fold) than that observed in direct lentivirus exposure of PHA/IL-2 treated PBMCs of the same donor (Supplementary Fig.2). The highest fold change (ca. 3-fold) in replication mediated by DCs was observed with the HIV-1 group M A74 virus (Supplementary Figure 2B). In using the MDDC and PBMCs of a different donor, we observed a similar level of enhancement of lentiviral replication with MDDC-mediated trans infection.

### Competitions of HIV and SIV isolates in human and chimpanzee PBMCs

Due to the inherent variation of culture conditions, logarithmic growth of viruses, and variability in viral monitoring assays, only gross differences in replication kinetic of different lentiviruses can be measured in monoinfection(45,55,56). Direct dual virus competitions at low multiplicity of infection in the same contained culture provides a relative, reproducible measure of replicative fitness with minimal variance(42,44,45,55,57). Thus, we performed pairwise competitions with 21 lentiviruses in both human and chimpanzee PBMCs using viruses (schematically illustrated in Figure 3A). A total of 253 separate pairwise competitions in *Hs*PBMC at an MOI of 0.005 were harvested after 12 days post infection, i.e. peak virus production. Dual infections at this MOI and in this time frame limits recombination to <0.1% of the observed dual virus replication(44,58,59). We obtained a limited supply of blood/PBMCs from *P.t. verus* (*Ptv*) to perform 82 pairwise competitions involving inter-lentiviral pairs (e.g. HIV-1 M strain versus SIVcpz rather than HIV-1 M subtype A versus HIV-1 M subtype B). *Ptv* was selected for PBMCs since this chimpanzee subspecies lacked infection by SIVcpz (as tested to date)(24–27). For inter-group/type competitions performed in both *Hs* and *Ptv*PBMCs, the extracted DNA was PCR amplified in a *gag* coding region, that is conserved among primate lentiviruses. Extracted DNA from intra HIV-1 group M competitions were amplified using the envelope primers described previously(42,44,55,57,60,61) (Figure 3A). PCR products were barcoded *and* sequenced by next generation deep sequencing. We then processed resulting reads through the *SeekDeep* pipeline which performs *de novo* clustering at a single base resolution(62). *SeekDeep* is advantageous as resulting measures are centered around the most abundant viral clones in the quasispecies/swarm and thereby avoid biases that may be engendered by direct mapping to a singular strain reference sequence.

**Figure 3:**
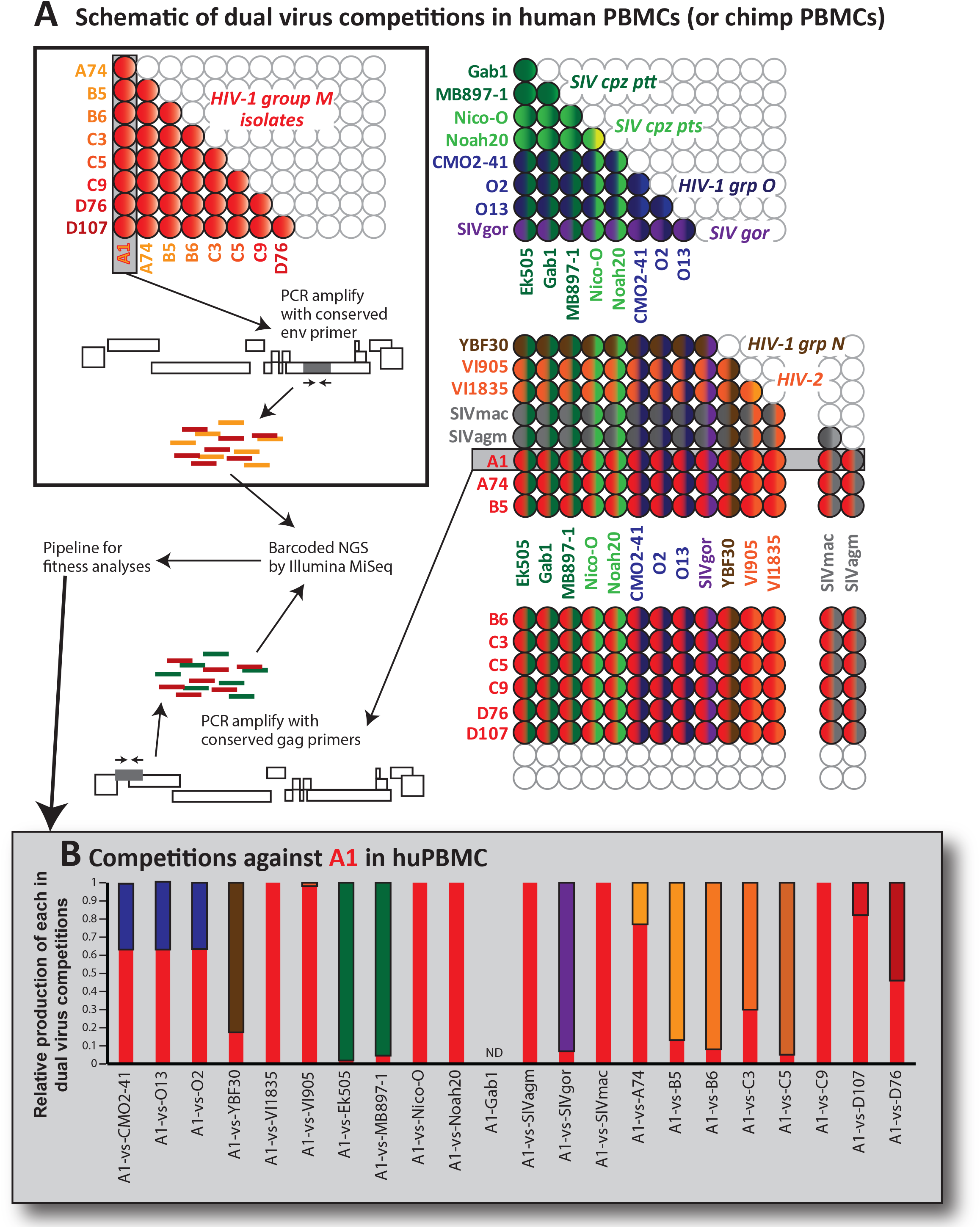
A) Schematic llustration of dual virus competitions in human and chimpanzee PBMC, (B) Results of the pairwise infections involving competitions of the HIV-1 group M A1 strain against all other primate lentiviruses. Relative production of both viruses as measure by NGS and SeekDeep is shown.

Supplementary figure 3 provides the detailed results from 335 individual competitions involving each pair of primate lentiviruses in human and *Ptv*PBMCs. Figure 4 provides the competition results of each primate lentivirus of different groups/types against HIV-1 group M isolates (Figure 4A and C) or SIVcpz isolates (4B and D) in *Hs*PBMCs (4A and B) and *Ptv*PBMCs (4C and D). In direct competitions, the human lentiviruses HIV-1 group O and HIV-2 were significantly less fit than SIVcpz and HIV-1 group M isolates in either *Hs*PBMCs (Fig. 4A) or *ptv*PBMCs (Fig. 4C) (ANOVA; Supplementary tables). Interestingly, SIVgor CP684 and SIVcpz strains, the progenitors of HIV in humans, could outcompete most HIV-1 group M isolates in *Hs*PBMCs (Fig. 4A) whereas HIV-1 and its progenitor, SIVsmm has the lowest replicative fitness. We were unable to propagate sufficient HIV-1 P RBF168 for this study but the SIVgor CP684 was over 90% identical in Gag, Pol and Env sequence to groups O and P(16), very similar to the homology between HIV-1 groups M and N and their respective SIVcpz progenitor.

**Figure 4:**
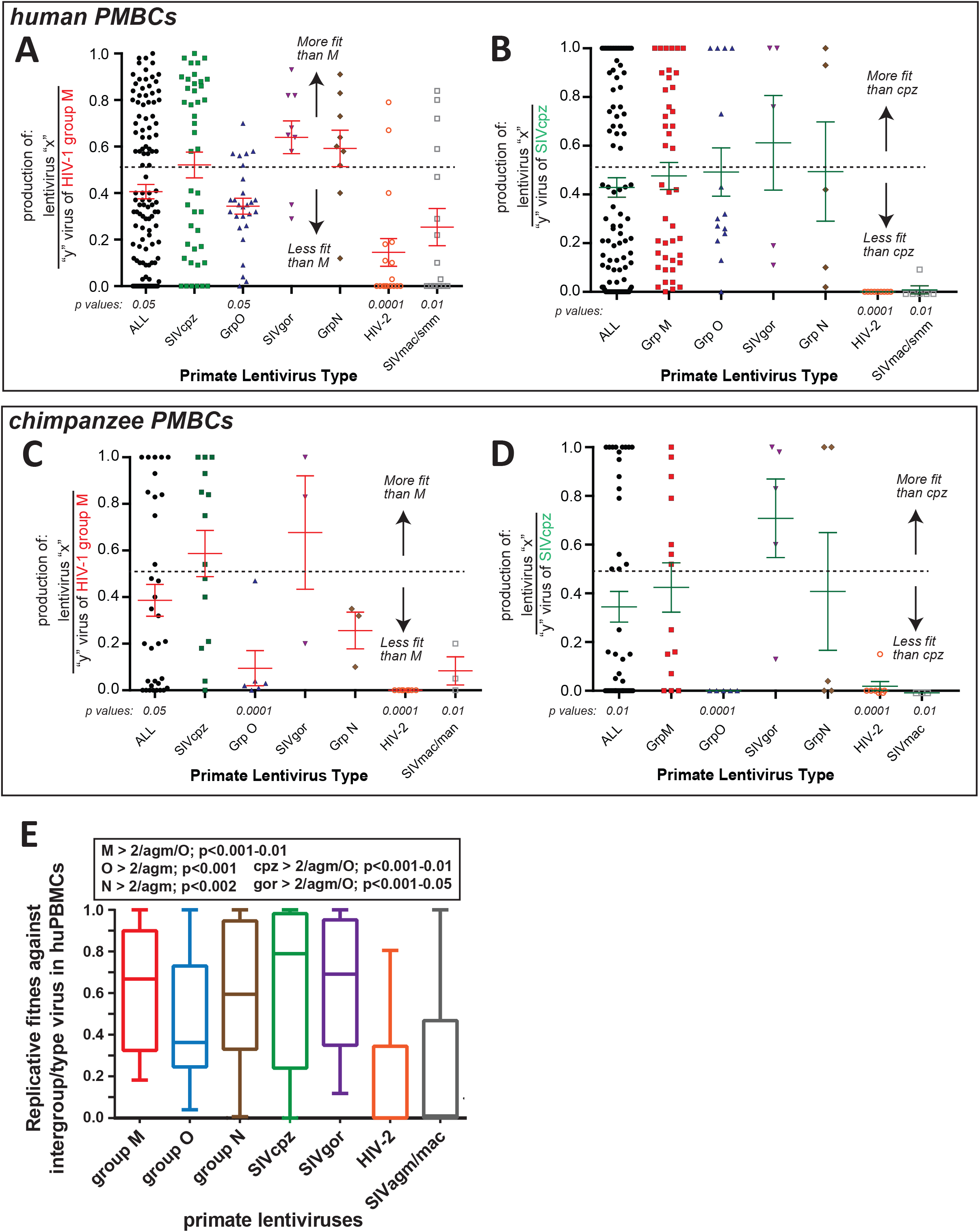
Fitness analyses of HIV-1, −2, and SIV isolates in human and chimpanzee PBMCs. As described in Figure 3, pairwise competitions were performed in human PBMCs and subset in chimpanzee PBMCs. The production from these dual virus competitions were PCR amplified, subject to NGS, and analyzed by SeekDeep to obtain the relative production of each virus in each dual virus competition. These values are plotted as the produciton of any lentivirus relative to the group M (A and C) or SIVcpz (B and D) in *Hs*PBMCs (A and B) or *Ptv*PBMCs (C and D). E. For this comparison, the replicative fitness of members of one primate lentivirus group/type (e.g. HIV-1 group M) competed against members of all other groups/-types was analyzed for an intergroup/type replicative fitness. This analyses excludes all of the intragroup/type dual virus competitions (e.g. HIV-1 group M A74 versus B5). The geometric mean and box of 95% of the relative fitness values with outlier whiskers is plotted for each group/type of primate lentivirus. Full statistical analyzes of all panels is provided in Supplementary tables 1 to 5, respectively.

In the competitions involving SIVcpz and HIV-1 group M in *Hs*PBMCs (Figure 4A and C), and the reciprocal plot in Figure 4B and D in *Ptv*PBMCs, SIVcpz strains trended to slightly higher replicative fitness than HIV-1 group M isolates. Again, the other SIV strains (not infecting chimpanzees) or the HIV-2 strains (originating from SIVsmm) were less fit than SIVcpz or group M HIV-1 in both *Hs* and *Ptv*PBMCs. HIV-2 or SIVsmm have not been identified in chimpanzees. Despite lower replicative fitness of HIV-1 group O versus M (Figure 4A&C), SIVcpz was only more fit than HIV-1 group O in *Ptv*PBMCs (Figure 4B) but not in *Hs*PBMCs (Figure 4D). The path of zoonotic jumps responsible for the origin of HIV-1 group O is not well defined and may involve an intermediate jump through chimpanzees(16,63).

The competition results are also presented as the replicative fitness values of each primate lentivirus within a group/type competed against the primate lentiviruses outside of this group/type in *Hs*PBMCs (e.g. HIV-1 group M isolates competed against HIV-1 O, N, HIV-2, SIVcpz, gor, agm, and mac; first bar in Figure 4E). HIV-1 groups M, N, SIVcpz and gor are significantly more fit than HIV-1 group O, HIV-2, SIVagm and mac (Figure 4E; ANOVA analyses in Supplementary tables).

### Replicative fitness differences among SIVcpz strains derived different chimpanzee subspecies

When comparing the replicative fitness of SIVcpz versus HIV-1 group M in *Hs*PBMCs and *Ptv*PBMCs, there were SIVcpz strains/viruses with high and low fitness. The competitions included SIVcpz_*ptt* isolated from the *Pan troglodytes troglodytes* subspecies and SIVcpz_*pts* found in the *P. troglodytes schweinfurthii*. SIVcpz_*ptt* strains were significantly more fit than SIVcpz_*pts* compared to HIV-1 group M in both *Hs* (Figure 5A) and *Ptv*PBMCs (Figure 5B) (ANOVA; Supplementary tables). The SIVcpz_*ptt* strains, EK505 and MB897 could equally compete with the more fit group M isolates, A1, A74, B5, and B6 whereas the SIVcpz_*pts* strain, Nico-0 and Noah-20 were out-competed in *Hs*PBMCs (Figure 5C). Gab1, the least fit SIVcpz_*ptt* strain was outcompeted by group M strains of high fitness but like all the SIVcpz strains, could compete with the HIV-1 group M isolates of the lowest replicative fitness (Figure 5D). It is important to stress that SIVcpz strains appeared most adapt at replicating in *Hs*PBMCs as compared to SIVsmm, SIVmac, and SIVagm, the latter of which replicated with slow kinetics on *Hs*PBMCs or susceptible primate cell lines (Figure 2 and 4). Among the SIVcpz strains, those isolated from *Pan troglodytes troglodytes* shared the closest genetic identity to group M (Figure 1) and were more adapted to replicating in human primary cells than SIVcpz from *P. troglodytes schweinfurthii*, a chimpanzee subspecies found primarily in West and East Africa and less in the Congo basin of Central Africa. Thus, the inherently higher replication fitness of SIVcpz_*ptt* versus SIVcpz_*pts* could have contributed to the cross-species transmission of the latter and evolution into HIV-1 Group M.

**Figure 5:**
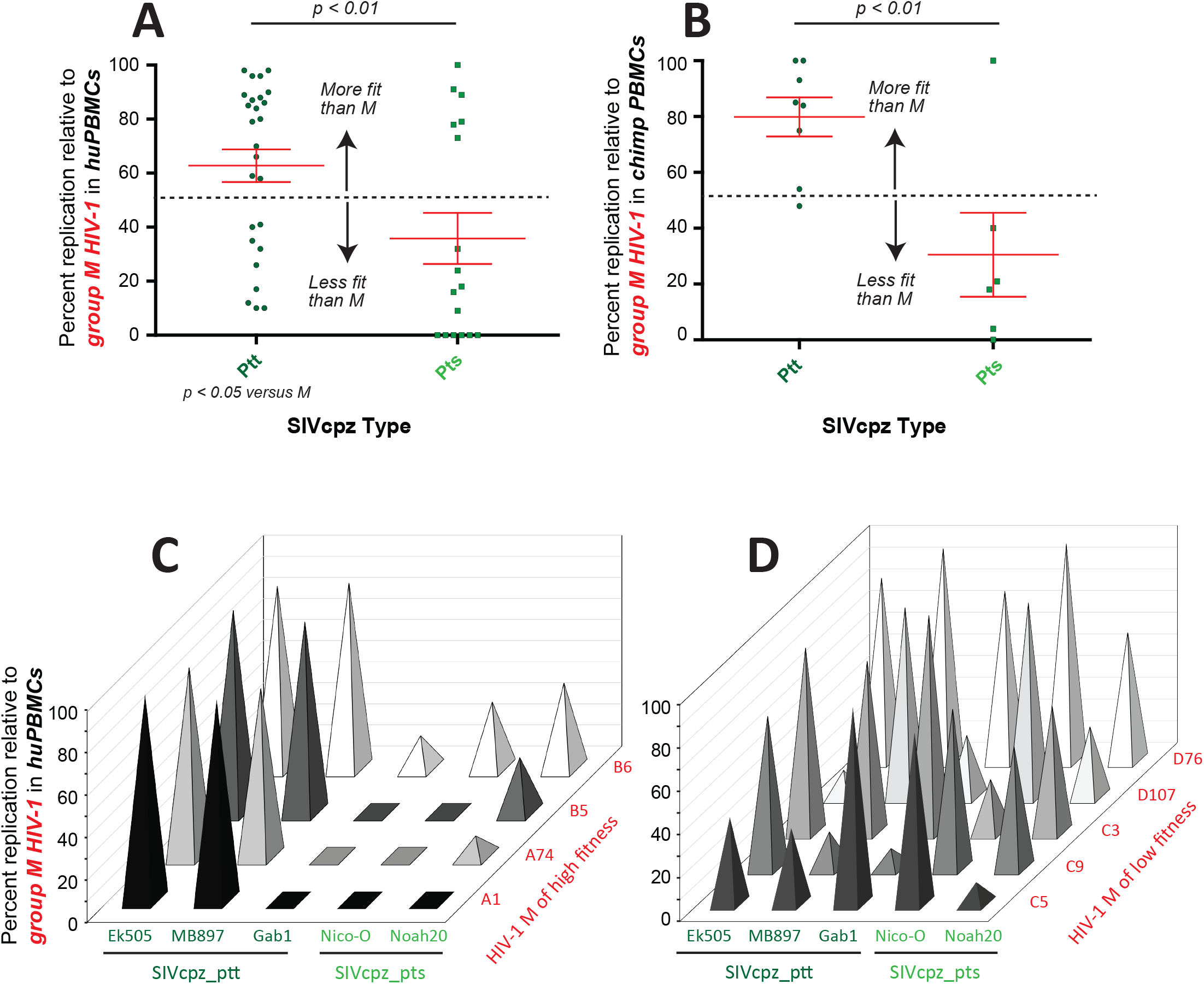
Fitness of various SIVcpz strains of *Ptt* and *Pts* competed against HIV-1 group M strains in A) *Hs*PBMC and B) *Ptv*PBMC. SIVcpz_ptt and _pts strains competed in human PBMCs against C) HIV-1 group M of high fitness and D) group M of low fitness.

### Impact of human restriction factors on primate lentivirus replication

Host factors such as Trim5a, APOBEC 3F/G/H, tetherin/BST2, and SamHD1 have all been shown to restrict xenotropic retrovirus replication and most have been associated with preventing cross species transmission of primate lentiviruses(29,64). We explored the ability of these restriction factors in human cells to reduce replication kinetics of NHP lentiviruses as one mechanism to prevent a jump into humans. For these studies, we utilized the cell line (U87.CD4.*Hs*CCR5) most supportive of HIV-1 group M replication and most restrictive to NHP lentiviral replication (Figure 2). In cells treated with siRNAs reducing the endogenous mRNA of Trim5α, tetherin/BST2, APOBEC 3F and 3G (by 80-90% for 5 days), we observed release of restriction and increase in lentivirus replication (Figure 6 and Supplementary Fig. 4). Reduction of human APOBEC3F and 3G levels resulted in minimal increases in lentiviral replication over 7 days (Supplementary Fig.4C and D), which is expected based on this restriction requiring an accumulation of deleterious mutations. Human Trim5α results in low level inhibition of all primate lentiviruses which is released with siRNA treatment (Supplementary Figure 5A). Trim5α is a stronger NHP restriction factor to prevent cross-species replication of lentiviruses. The strongest restriction of NHP lentiviruses in these human cell lines appears to be mediated by tetherin/BST2 (Figure 6 and Supplementary Fig. 4). SIVmac and SIVman replication increased by 5-6 fold with a >80% knockdown of tetherin mRNA over a 9 day infection. Tetherin inhibits even HIV-1 group M and O isolates at a low level. In contrast, there was no apparent inhibition of SIVcpz strains by human tetherin.

**Figure 6.**
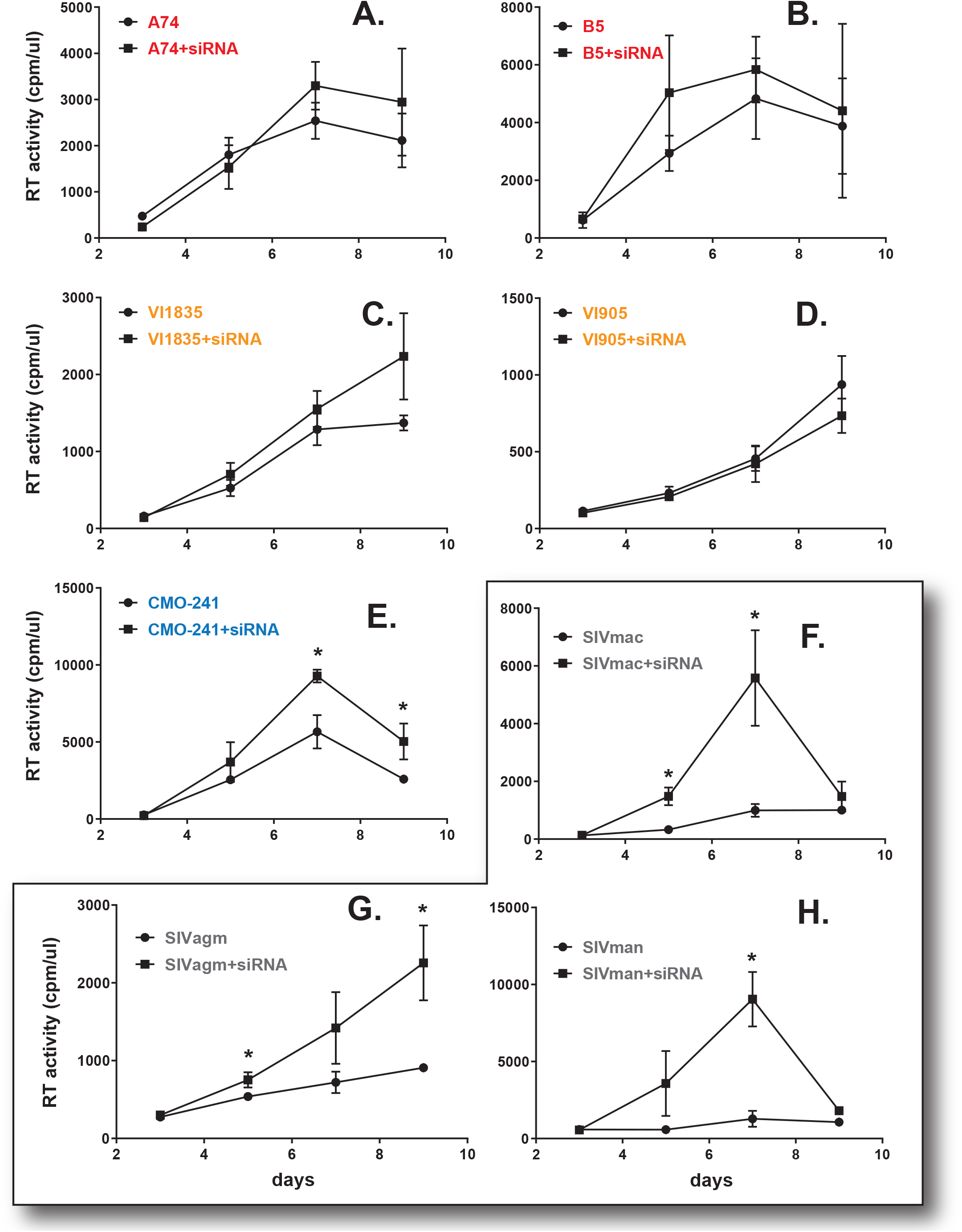
Primate lentivirus replication in human CD4+ U87 cells expressing HsCCR5 and pre-treated with siRNA to Tetherin/BST2. HIV-1 group M A74 (A), B5 (B), HIV-2 VI1835 (C), VI905 (D), HIV-1 group O CMO-241 (E), and SIVmac (F), agm (G), and man (H) were used to infected CD4/HsCCR5 U87 cells treated with scrambled siRNA or siRNA specific to mRNA Tetherin/BST2. All infections and treatments were performed in triplicate. * represents p < 0.05, multiple one-tailed T tests.

Based on the pairwise virus competitions with the 21 lentiviruses performed in HsPBMCs, we calculated the average replicative fitness value of each lentivirus based on only the intergroup/type competitions and removing all intragroup/type fitness values (excluding e.g. group M A74 versus group M B5). As described in Supplementary Figure 5, the average intergroup/type fitness was consistent with replicative fitness derived from every competition against an HIV-1 group M or SIVcpz isolate as described in Figure 3. The average intragroup/type fitness values of each of the 21 lentiviruses were then compared to the relative restriction by human Trim 5α (i), Tetherin (ii), APOBEC 3F (iii), and 3G (iv) and we observed no significant correlations (Figure 7A). However, when the average fitness of only SIVs were plotted, a significant inverse correlation was observed between replicative fitness and release of restriction by Trim 5α, Tetherin, or the additive effects by all four factors (Figure 7Bi, ii, and v). Again, the SIVcpz strains (especially cpz_*Ptt*) had the highest replicative fitness and appeared to be the least restricted in human cells.

**Figure 7:**
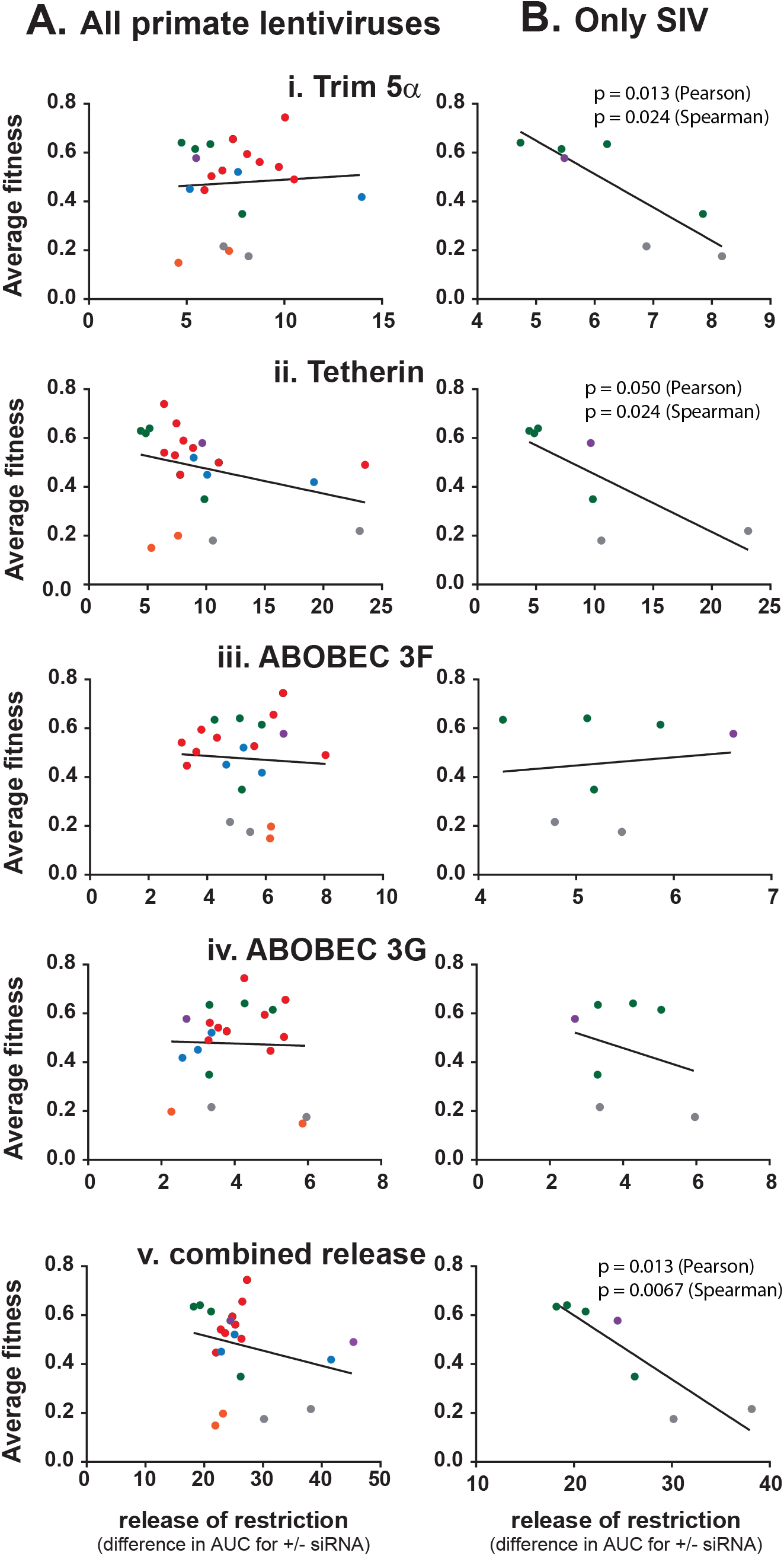
Comparing the average replicative fitness to the release of restriction (mediated by siRNAs). The average replicative fitness for each primate lentivirus was determined from the pairwise competitions in HsPB-MCs as described in Supplementary Figure 3. Each primate lentivirus was also used to infect cells in the presence of a specific siRNA to restriction factor or a scrambled siRNA. The area under the curve for the 9 day infection with specific siRNA was subtracted from that with scrambled siRNA. (A) For each primate lentivurs, plots of average replicative fitness versus the relative release of replication due to Trim 5a (i), Tetherin/BST2 (ii), ABOBEC 3G (iii), 3F (iv), and all four combined (v). (B) The same plots as in (A) but without the human lentiviruses.

### Cytopathic phenotype of HIV-1, HIV-2 and SIV in ex-vivo lymphoid tissues

Unlike PBMCs, human tonsils are permissive cells that do not require stimulation with mitogens or IL-2 for HIV propagation(65). To determine the cytopathic effect and CD4 T-cell depletion rate by these primate lentiviruses, human tonsils were infected with the same infectious titers and monitored for virus production and CD4 T-cell depletion based on pyroptosis and/or necrosis relative to uninfected controls(53,54). It is important to note that depletion of CD4 T-cell in human tonsils by R5-using HIV-1 subtype B strains (average of 10% herein with the HIV-1 group M isolates) is lower than reported with X4-using HIV-1 strains, typically reaching a 65% loss in tonsillar CD4+ T cells(54). In this study, the highest CD4+ T cell depletion was observed with SIVmac (average 27%), followed by A74 (18%), VI1835 (16%) and D107 (12%) (Figure 8A). Despite the relatively low replication in CD4+ T-cells, HIV-2 and SIVmac/agm still depleted significant levels of CD4 T-cells in human tonsils (Figure 8B). Exposure to the human tonsillar tissue to SIVcpz strains (*Ptt* and *Pts*) resulted in minimal CD4+ T cell depletion (1 to 3.5%; Figure 8A) despite high levels of replication in these tissues (Figure 8B). SIVcpz_*Pts* Nico had the highest ratio of virus production versus CD4+ T-cell depletion in the tonsillar tissues. All the primate lentiviruses within HIV-1 groups (M, N, O), HIV-2, SIVgor and SIVagm/mac were significantly better at depleting CD4 T-cells than SIVcpz (except group N and SIVgor, limited data points). In contrast, there was no difference in the levels of CD4 T-cell depletion in the tonsillar tissues between any of the types/groups when SIVcpz was not included in the ANOVA analyses.

**Figure 8:**
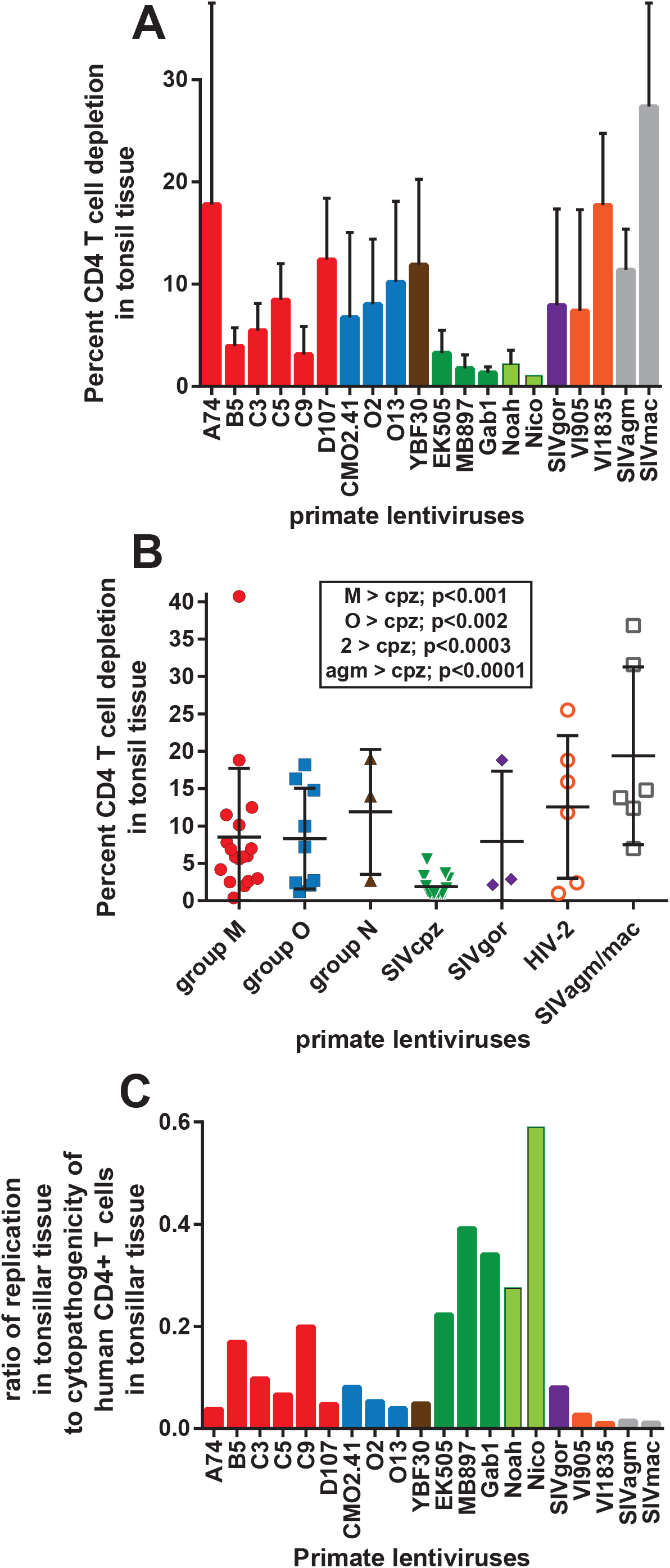
Primate lentiviral replication in tonsillar tissue and huPBMC fitness (A) Percentage of T-cell depletion; (B) CD4 T-cell depletion levels within a group or type of primate lentivirus; (C) Comparing the replication in tonsillar tissue to cytopathogenicity

No correlation was observed between the relative replication in the tonsillar tissue and the level of CD4 depletion by these viruses. SIVcpz had the highest replicative fitness in human PBMCs and in the tonsillar tissues. In contrast, SIVcpz infection was associated with minimal cytopathogenicity in human tonsillar tissue resulting in the greatest ratio of virus production to CD4+ T cell death of any primate lentivirus (Figure 8C). In contrast, the HIV-2 isolates, SIVagm, and SIVmac showed an average depletion of CD4+ T cells, i.e. comparable to that observed with HIV-1 group M, O, and all other primate lentiviruses (except SIVcpz) (Figure 8A and B). Based on the poor virus production in human tonsillar tissue (as well as average fitness in HsPBMCs), the ratio of replication to CD4 T cell depletion of HIV-2 and SIVagm/mac was the lowest for all primate lentiviruses (Figure 8C). As described below, these findings might suggest that pathogenicity/virulence could have evolved as a trait in humans following a zoonotic jump.

## Discussion

Phylogenetic studies have revealed a complex history of interspecies jumps of lentiviruses among catarrhine monkeys or “old world” monkeys. Based on the discovery of an endogenous primate lentivirus in lemurs of Madagascar, best estimates place the introduction of lentivirus into monkeys approximately 17 million years ago from a jump of a primordial lentivirus which then branched into FIV and SIV(66). Despite evidence of primate lentiviruses in old world monkeys for millions of years, our knowledge of this evolution is largely based on the jumps during the past 10,000 years resulting in the existing lentiviral lineages in primates, many of which are unstable in their current host species. An example of this instability is the jump of SIV from sooty managabeys into humans in 1930s(67) which resulted in HIV-2 with a peak epidemic of 1-2 million, limited transmission success, and the progressive loss in prevalence within humans, even before the introduction of antiretroviral treatment. HIV-1 group O, N, and P had even less success surviving in humans following introductions from the gorilla and/or chimpanzees; jumps dated to sometime in the 1920s for group O and 1960s for N(22,67,68). In contrast, HIV-1 group M emerged in humans with a jump from chimpanzees (*Ptt*) in the Congo basin after the turn of the 19^th^ century, an event that has now resulted in over 40 million deaths with another 37 million infected humans(67–69). There is no clear evidence of attenuation of HIV-1 virulence or reduction in transmission efficiency but some in vitro studies suggest this possibility(39,70).

This study was designed to compare the competitive replicative fitness and virulence of different lentiviral strains isolated from various primate species. Exposure of a primate to a xenotropic lentivirus is unlikely to result in a cross-species transmission event due in part to protective barriers, e.g. innate defenses and various restrictions factors. Nonetheless these restriction factor/innate defenses appear to show variable protection against different, foreign lentiviruses. To understand the overall restriction of primate lentiviruses in primary human cells susceptible to HIV, we compared the replicative fitness of primate lentiviruses in both human and chimpanzee PBMCs by performing over 250 with 21 lentiviral isolates of HIV-1 groups M, N, O, HIV-2, SIVcpz, gor, smm, and agm. To determine relative virulence upon a possible jump to humans, we compared the relative replicative fitness of these primate lentiviruses to the inhibition by various human restriction factors and the relative depletion of CD4+ T cells in human tonsillar tissue.

To summarize, SIV strains from *Ptt* and *Gor* and HIV-1 group N appeared slightly more fit than even HIV-1 group M replicating in both human and chimp PBMCs. However, as previously published, there was a wide range in replicative fitness among HIV-1 group M subtypes with subtype A and C strains again being consistently less fit(37,42,55,57). When these HIV-1 group M strains of lower fitness were removed from the analyses, there was no significant difference in the replicative fitness of SIV from chimpanzees and gorillas and the HIV-1 group M and group N isolates found in humans. SIVcpz or SIVgor (progenitor of HIV-1 group P) are significantly more fit than HIV-1 group O in human PBMCs. Finally, SIV from African green monkeys and Sooty mangabey had the lowest replicative fitness in both human and chimp PBMCs, similar to the low replicative fitness observed with HIV-2. We determined that the relative level of replication of these non-human primate lentiviruses in human and chimp PBMCs reflect the cumulative inhibition by the various human restriction factors, with tetherin/BST2 being the most restrictive. Based on the cumulative data presented herein, it is clear that SIVcpz and SIVgor have the greatest ability to replicate in human PBMCs. However, the SIVcpz derived from *Pts*, which did not establish lentivirus epidemic in humans, was less fit in human PBMCs than SIVcpz of *Ptt*. SIVcpz from *Pts* or *Ptt* were also inhibited the least by human restriction factors. Based on the similar overlap and invasion of humans into the *Pts* and *Ptt* habitats in the Rift Valley of East Africa, Congo Basin of Central Africa, and the tropical south of West Africa, it is unlikely that humans would be exposed less to *Pts* than to *Ptt* chimpanzees. Findings presented herein suggest that the SIVcpz of *Ptt* compared to SIVcpz of *Pts* was more compatible for cross species transmission to humans and gained the foothold necessary for transmission and human adaptation.

The ability of a lentivirus to replicate in the primary susceptible cells in a host does not always correlate with pathogenesis. A strong example is the lack of disease in African Green Monkeys despite the high viral loads of SIVagm produced by an abundance of infected T-cells in these animals(46,47). There has only been one study that has characterized disease progression of a chimpanzees infected by SIVcpz (a *Pts* strain) in the wild but raised in captivity. With this animal there was little evidence of disease despite an increasing viral load during the length of infection(48). In humans, several studies have described a direct correlation between the rate of disease progression and the replicative fitness of the infecting HIV-1 isolates in primary T-cells(37,44,45,71). This is best illustrated by the low replicative fitness of HIV-1 isolates derived from patients with slow or non-progressing disease versus the high replicative HIV fitness among rapid progressors. In this study, we compared the cytopathogenicity of these primate lentiviruses in primary human lymphoid tissue, i.e. resected tonsillar tissue. There was a clear disconnect between the ability of non-human primate lentiviruses to propagate in human tonsillar tissue and the pyroptosis/cell death of the CD4+ T cells within the same tissue. Similar levels of T-cell death was mediated by HIV-2, SIVagm, and SIVsmm with low replicative fitness than by HIV-1 group M/N and SIVgor with high replicative fitness. In contrast, SIVcpz having the highest replicative fitness in tonsillar tissue (as well as in human and chimp PBMCs) was associated with minimal pyroptosis and all manner of T-cell death in this tonsillar tissue. The inability of SIVcpz to mediate cytopathogenicity in human T-cells may correspond to the minimal pathogenesis or lack of disease described in most SIV infected chimpanzees(48,49). Again, in contrast to the most fit SIVcpz’s, SIVsmm (the progenitor of HIV-2) and HIV-2 were highly cytopathogenic in these tonsillar tissues despite having the lowest replicative capacity. These findings are consistent with slow disease progression associated with HIV-2 infected humans(51,52). HIV-1 group O and SIVgor (the progenitor of HIV-1 group O) had high cytopathogenicity and moderate replicative fitness in these tonsillar tissues but not as low as HIV-2/SIVsmm. HIV-1 group O infected humans progress faster to disease than those infected with HIV-2 but slower than those infected with HIV-1 group M(63).

The relative virulence of a virus in their host species is of course impacted by a multitude of host factors (e.g. innate and acquired immune responses, restriction factors). We propose that lentivirus virulence following cross-species transmission may relate to replicative fitness and per virus pathogenicity, both of which are influenced by the level of inhibition by innate restriction factors. This hypothesis appears to apply with the SIVgor and SIVsmm establishing the group O and HIV-2 spread in humans(2,22,68). Despite the close genetic relationship of SIVcpz_*Ptt* and HIV-1 group M, the SIVcpz strains from *Ptt* and *Pts* and HIV-1 group M strains had similar replicative fitness in human cells but the SIVcpz strains did not result in cytopathogenicity due to bystander effects or direct infection. This inconsistency between SIVcpz_*Ptt* and HIV-1 could relate to an adaptation/evolution of SIVcpz in humans to escape restriction factors/barriers following a jump. However, there is little evidence that SIVcpz is restricted from replication in human PBMCs as are other SIV lineages. It is feasible that an increased cytopathogenicity evolved early in the human epidemic during a period of high viral burden and efficient transmission. As HIV-1 group M diversified during the past 100 years, HIV-1 subtypes may have evolved with differences in virulence and transmission efficiency. Increase virulence of HIV-1 subtype D over subtypes A and C has been well documented within infected cohorts and using similar *ex vivo* studies described herein(37,40,42,55,57). It is also feasible that the HIV-1 epidemic relates to the cross-species transmission of a rare SIVcpz strain resulting in exceptionally high virulence (replicative fitness + cytopathogenicity) in humans. The chimpanzees carrying this SIVcpz may now be extinct. The rarity of this SIVcpz strain in chimpanzees might explain why a lentiviral epidemic did not appear earlier in human history considering the clear interaction of chimpanzees and humans in central Africa. With this scenario, the HIV subtypes may have evolved with differential reductions in virulence and transmission efficiencies in humans. Clearly, neither of these hypotheses can be proven without the availability of human lentiviral samples closer to the cross-specific transmission event responsible for the HIV-1 epidemic.

## Materials and Methods

### Cells

#### Primary and Cell lines

HIV negative subjects provided whole blood from which peripheral blood mononuclear cells (PBMC) were separated using the Ficoll-Hypaque density centrifugation technique. Whole blood from Chimpanzee was obtained from an HIV negative Chimpanzee (*Pan troglodytes verus*) from the Texas Primate Facility and shipped overnight to Case Western Reserve University, Cleveland and PBMCs were separated as described above. All PBMCs were stimulated for 3-4 days with 2ug/ml of phytohemagglutinin (PHA; Gibco BRL) and 1 ug/ml of interleukin-2 (IL-2; Gibco BRL) in complete RPMI 1640 containing 2mM L-glutamine. U87 human glioma cells expressing CD4 and CCR5 were obtained through the AIDS Research and Reference Reagent Program and in maintained in DMEM supplemented with 10% FBS, Fetal Bovine Serum (FBS; Mediatech, Inc.), Penicillin (100 U/ml), Streptomycin (100 ug/ml), Puromycin (1ug/ml) and G418 (300 μg/ml). 293 T-cells (human embryonic kidney cells) were grown also grown in DMEM media.

#### Human tonsils

Human tonsils removed during routine tonsillectomy (provided by the Human Tissue Procurement Facility of the University Hospitals Case Medical Center Hospital, Cleveland, Ohio) was received within 5 hrs of excision. Use of anonymous surgical waste for experimental HIV infections was approved by the IRB under UHIRB 01-02-45. To prepare human lymphocyte aggregate cultures (HLACs), we removed the cauterized tissue from the tonsil as well as the capsule and the fatty parts surrounding the tonsil. Bloody or inflamed parts were also discarded from the tissue. The tonsil tissue was dissected into small pieces by hand with a surgical scissors and the pieces were squeezed using a flat surface. Cells were dispersed and washed in Phosphate-Buffered Saline (PBS) and passed through a 70μm cell strainer (BD Falcon). Finally, isolated HLACs were plated in 96-well U-bottomed plates (Corning, Inc.) at a concentration of 2×10^6^ cells per well in 200μl RPMI 1640 media containing 10% FCS, 100 U/mL penicillin, 100 μg/mL streptomycin sulfate, 2.5 μg/ml fungizone, 2 mM L-glutamine, 10 mM HEPES, 1 mM sodium pyruvate and 1% non-essential amino acids mixture. One day after HLACs preparation, cells were inoculated with virus in triplicate (0.002 multiplicity of infection; (MOI) per 2×10^6^ cells per well in 200 μl). After overnight infection, cells were washed, and supernatants harvested at days 4, 7, and 10 post infection without dispersing the pellet. Lentiviral replication was assessed using an RT assay as described previously(72).

#### Monocyte-derived dendritic Cells (MDDCs)

MDDCs were isolated as described previously(73). Briefly, CD14^+^ beads (Miltenyi Biotech) were used to isolate CD14^+^ monocytes from healthy donor human PBMCs. These were then cultured in the typical 10% FBS/RPMI media and supplemented with 10ng/ml IL-4 and 50ng/ml GM-CSF (Miltenyi Biotech). MDDCs were matured by overnight stimulation with 100ng/ml LPS (Sigma) 5-6 days after initiation of the cultures. Mature MDDCs were pelleted, washed and resuspended in fresh medium ready for co-culture with PBMCs and infection with viruses.

### Viruses

Twenty six primate lentiviruses, primary isolates or molecular clones were used for this study and included HIV-1 group M (n=10), N (n=1) and O (n=3), and HIV-2 (n=2) and from non-human primates; SIVcpz (n=6), SIVgor (n=1), SIVagm (n=1), smm (n=1), and mac (n=1). A full list of the viruses, their origins, phylogenetic relations in env and gag, titers are outlined in Figure 1A and B. The HIV-1 group M and group O isolates have been used previously(42) and most were originally obtained from the AIDS Research and Reference Reagent Program of the National Institute of Health. The well characterized group molecular O clone CMO2.41 was also used as one of the representatives of HIV-1 group O(74). SIVgor (CP684) and three of the SIVcpz strains (EK505, Gab1-1, MB897-1) were gifts from Beatrice Hahn. The remaining SIVcpz and SIVagm, smm, and mac isolates were provided by the Institute of Tropical Medicine in Antwerp, Belgium. The SIVcpz strains Nico and Noah were obtained from a chimpanzee infected in the wild, maintained in captivity, and whose virus was then used to infect another chimpanzee(48).

Plasmids of the proviral clones (CMO2.4, CP684, EK505, Gab1-1, MB897-1) were used to generate infectious virus by transfecting 293T cells using the Effectene transfection reagent (QIAGEN, Valencia, CA). Supernatant containing virus was collected 2 days post transfection, clarified by centrifugation at 2,500 rpm for 10 mins and purified through a 0.45um pore-size filter (Millipore, Billerica, MA). Transfection derived supernatants were subsequently briefly passaged in either human PBMC while all other primary isolates were briefly passaged in PBMCs to generate virus stocks. The tissue culture infectious dose for 50% infectivity (TCID_50_ expressed as IU/ml)) of propagated viruses was calculated by the Reed and Muench method as described previously(42). Virus productions in supernatant was determined by a positive RT activity (two standard deviations above the negative control). RT activity is the preferred assay considering the enzyme is present in all HIV or SIV strains.

### Assessment of CD4^+^ T-cell depletion and RT activity in human tonsil histocultures

Twelve days post infection, HLACs were analyzed by fluorescence-activated cell sorting (FACS). Uninfected and infected HLACs were stained with Live/dead^®^ violet (Invitrogen, Grand Island, NY), Annexin V-APC (e-biosciences, San Diego, CA) and with cell surface markers α-human CD4 PE (Biolegend, San Diego, CA), α-human CD3 PerCP (BD Biosciences) and α-humanCD8 FITC (BD Pharmigen). Cells were acquired in a LSRII flow cytometer (BD Biosciences, San Jose, CA) and data analyzed with FloJo software (Tree Star, Inc. Ashland, OR). Supernatants from each tonsil infection was also assayed for RT activity(72).

### Ex-vivo growth competition assay

*Ex vivo* pathogenic fitness for the isolates was determined by infecting human and chimpanzee PBMCs using methods described previously(42,44,55,57,60,61). In human PBMC, pathogenic fitness of HIV/SIV isolates was performed by full pair wise competition using 9 group M (2 each of subtypes A, B, D; and 3 C), 3 group O, 1 group N, 5 SIVcpz (3 from *Ptt* and 2 from *Pts*), 2 HIV-2 and 1 each of SIV*smm*, *agm* and *mac*. For chimpanzee PBMC the competition was restricted to a selected number of isolates due to the limitation of cell number. All competitions were performed in 48 well plate containing 2×10^5^ PBMCs and equal multiplicity of infection (0.005) for both viruses, i.e. 1000 IUs. Virus production was monitored by measuring the RT activity until peak production at day 12 when cells were harvested and stored at −80°C.

### PCR and sequence analyses

Due to the closer genetic relationship between HIV-1 group M, DNA extracted from intra-HIV-1 group M competitions were PCR amplified using envelope primers described previously(42,44,55,57,60,61). Briefly, C2-C3 region of envelope was amplified by PCR using the primers E80 and E125 primers(44). For the DNA extracts of the competitions using the more divergent HIV-1/HIV-2/SIV viruses, conserved primers that spanned the primer binding site (PBS) PBSdt-F AAAATCTCTAGCAGTGGCGCCCGAACAG (position 622-649 in HxB2; 806-835 in SMM239) and the 5’ end of gag (MAp17) GAGdt-R TTTCCAGCTCCCTGCTTGCCCATACTA (position 890-916 in HxB2 and 1153-1179 in SMM239) was employed for PCR. The NGS protocol to amplify and sequence the 300 competitions using 454 and MiSeq has been recently described. It is important to stress that the same results (+/−2% variance) was observed with both NGS platforms despite the increased error rates with 454. The alignments and analysis pipeline described below “count” the number of reads that cluster with one versus the other lentiviruses in the competition. All quasispecies of lentiviruses are genetically distinct using this amplified and sequenced gag or env region.

### Competition Sequence Data extraction and Analyses

Sequence data was extracted and analyzed using *SeekDeep*, a software suit that enables *de novo* clustering thereby avoiding potential artifacts that may occur due to alignment directly to reference sequences in situations of divergent sequence populations. It provides start-to-finish workflow from raw sequences files to population-level clustering, tabular and graphical summaries(62). The program has three components namely: extractor, qluster and process Clusters.

Sequence reads were first separated at the sample level based on their MIDs/bar codes followed by the detection and removal of forward and reverse PCR primers to leave the targeted region. Additional filtering was performed to ensure expected target length and to remove low quality reads as indicated by low quality or ambiguous (N) bases in the sequences. Next, was the clustering phase which involved the collapsing of unique reads except for high quality differences. Clusters that were up to 2 bases different from a more abundant haplotype were collapsed. Clusters composed of a single read were removed. During the *SeekDeep* process clustering phase, a final filtering was performed to remove low abundance artifacts and chimeric haplotypes, determined as a haplotype that was a combination of two other more abundant haplotype clusters (>2-fold relative to potential chimera), low frequency chimeric haplotypes that represented combinations of and low abundance artifacts. These final haplotype clusters were than mapped to known input strain sequences allowing up to 5 mismatches and renamed appropriately. The relative abundance of the strains in each competition was then calculated and interactive graphs of the results were created to assess quality and view the data(62).

### Phylogenetic and Evolutionary analysis by Maximum Likelihood method

All group M and O strains used in the study had been partially sequenced in the env gene (C2-V3; 480 nucleotides) and gag (MA p17; 350 nucleotides) as described previously (Abraha et al 2009). The group N and some of the SIV strains are available as full or partial genome sequences in the HIV Los Alamos database. Sequences were aligned using Clustal X incorporated in the MEGA software. Reference sequences of various HIV types and groups were also included. The evolutionary history was inferred by using the Maximum Likelihood method and Kimura 2-parameter model(75). Initial tree(s) for the heuristic search were obtained automatically by applying Neighbor-Joining and BioNJ algorithms to a matrix of pairwise distances estimated using the Maximum Composite Likelihood (MCL) approach, and then selecting the topology with superior log likelihood value. The tree was drawn to scale, with branch lengths measured in the number of substitutions per site. Evolutionary analyses were conducted in MEGA X(76).

### SiRNA inhibition of restriction factors

U87-CCR5 cells were seeded with 5×10^4^ per well in DMEM and incubated in 48 well plates overnight at 37°C. On the next day, cells were washed with PBS and DMEM media replaced with 210ul of Opti-MEM (Thermofisher). siRNA against Trim 5α, APOBEG 3F and G, and Tetherin obtained from Santa Cruz Biotechnology Inc. was dissolved respectively and resuspended in an accompanying diluent. Cells were then transfected with siRNA (10nM final concentration) using Lipofectamin RNAi Max reagent (Invitrogen) as follows: 0.6ul Lipofectamin in 20ul Opti-MEM and 0.25ul of the respective siRNA in 20ul Opti-MEM as described previously(77). These were then mixed and added drop wise in each well and incubated for 36 hrs. Cells were then washed with PBS, diluted virus added at a MOI of 0.001 and incubated at 37°C for 5 hrs in DMEM. Cells were washed and resuspended in 400ul DMEM and supernatants were harvested on days 3, 5, 7 and 9 post infection and tested for RT activity. All experiments were done in triplicates; and included virus+scrambled siRNA, virus+siRNA, as well as negative controls (siRNA alone and cells alone). Endogenous expression of these restriction factors is rarely detected by Western blots using antibodies. The reduction of the restriction factor mRNA levels with these optimized siRNA was between 65-95%. Please note the same U87 cell culture treated with one specific siRNA was split and utilized for infections with all lentiviruses.

### Statistical analyses

We employed the spearman correlation to determine if there was any relationship between the genetic distance and the fitness in human PBMC, as well as fitness and CD4^+^ T-cell depletion in the tonsil by the isolates.

## Acknowledgements

We thank Drs. Beatrice Hahn, Jonathan Heeney, and Brandon Keele for providing primate lentiviruses for this study. Support for this study to E.J.A. was provided by the NIH/NIAID R01 AI49170 and CIHR project grant 385787. Infrastructure support was provided by the NIH CFAR AI36219 and Canadian CFI/Ontario ORF 36287. Efforts of J.A.B. and N.J.H. was provided by NIH AI099473 for and AI099473. D.H.C. was funded by VA and NIH AI AI080313 for this study.

---

**Supplementary Figure 1:**
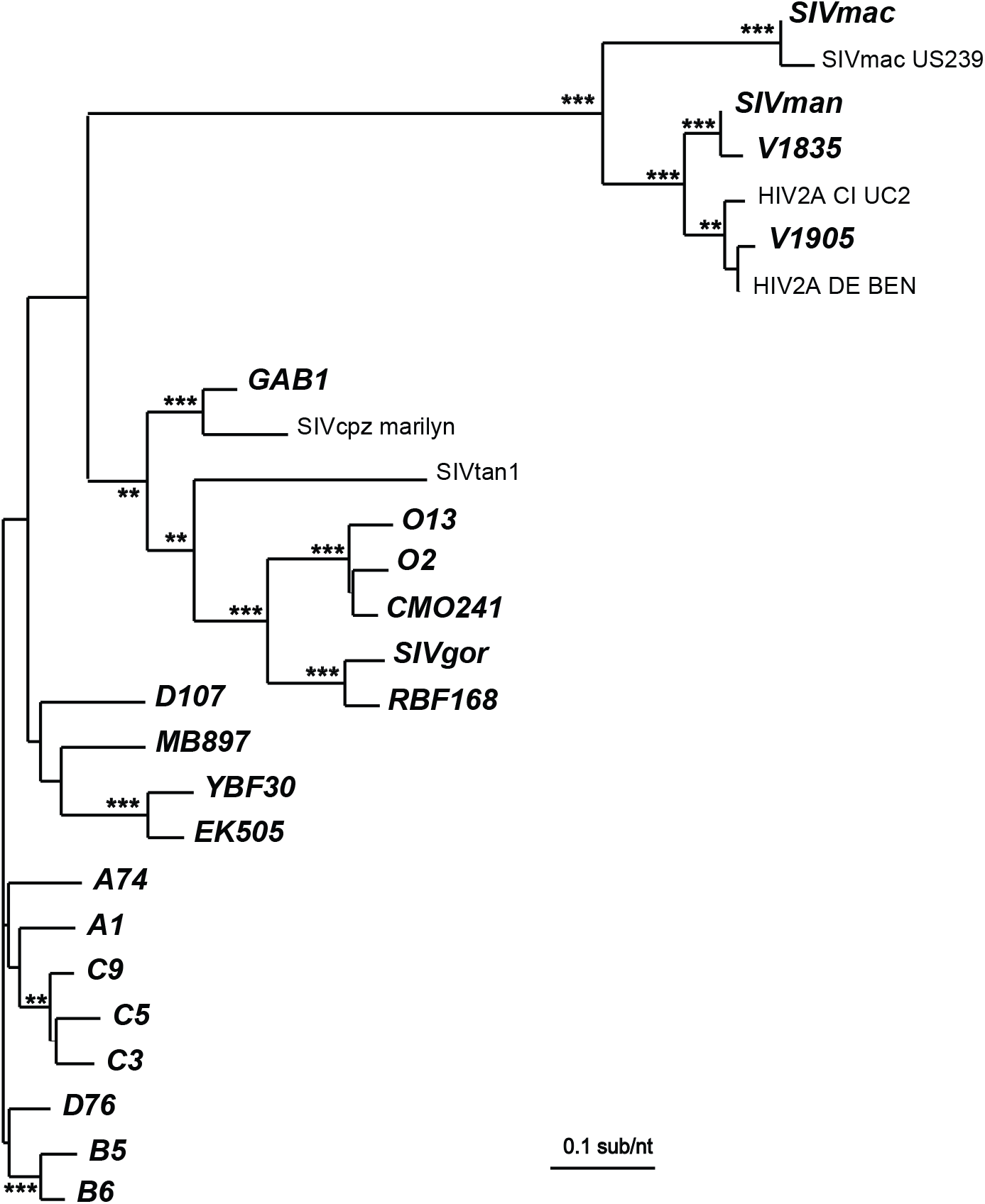
Neighbor joining phylogenetic tree of the *gag p24* sequences of primate lentiviruses with reference strains.

**Supplementary Figure 2:**
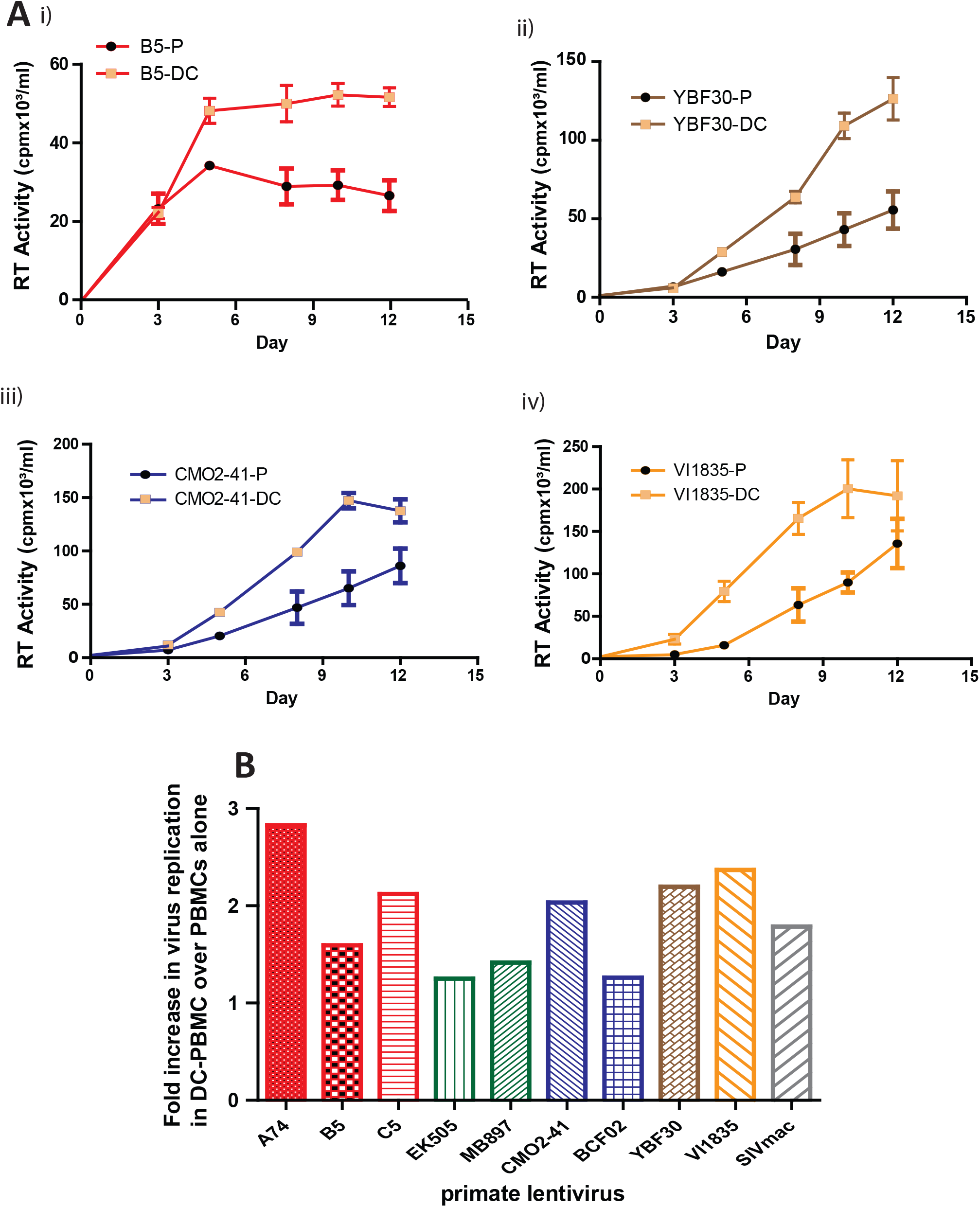
Enhancement of HIV and SIV replication in PBMC+dendritic cell cultures. A) Comparison of the replication kinetics of some isolates in PBMC (P, black filled circles) and PBMC+DC (DC, gold filled squares) co-cultures. B) Fold difference showing enhancement in replication by DCs.

**Supplementary Figure 3a:**
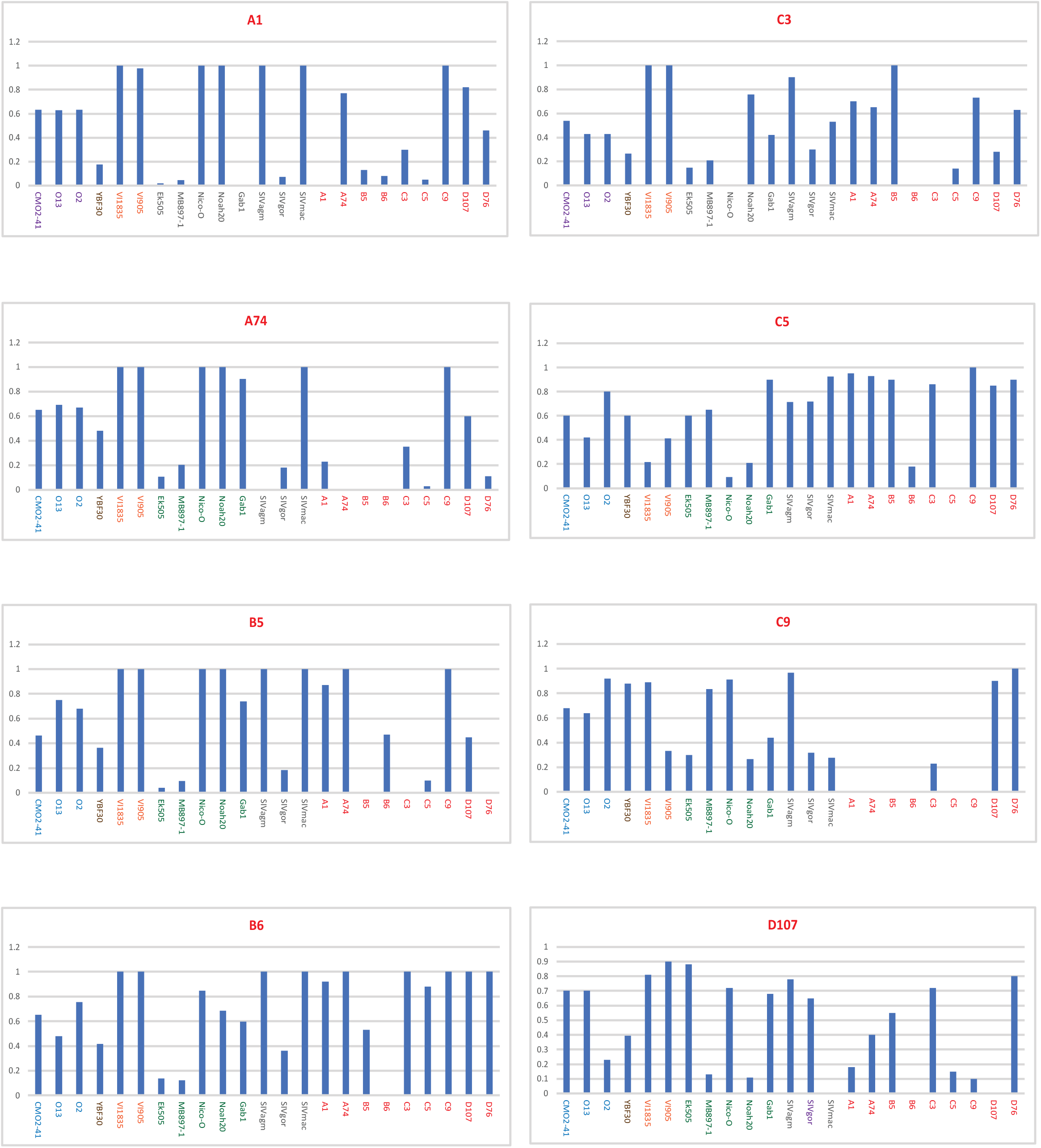
Fitness of primate lentiviruses in human PBMC. Primate lentivirus were added to *Hs*PBMCs for a full pairwise competition. Production of the “titled” virus in each chart is presented as fraction of the total when competed against the virus on the X axis. Relative virus production was measured from PCR and deep sequencing analyses using the SeekDeep pipeline. All competitions were performed in duplicate with a 10% observed variance in fitness.

**Supplementary Figure 3b:**
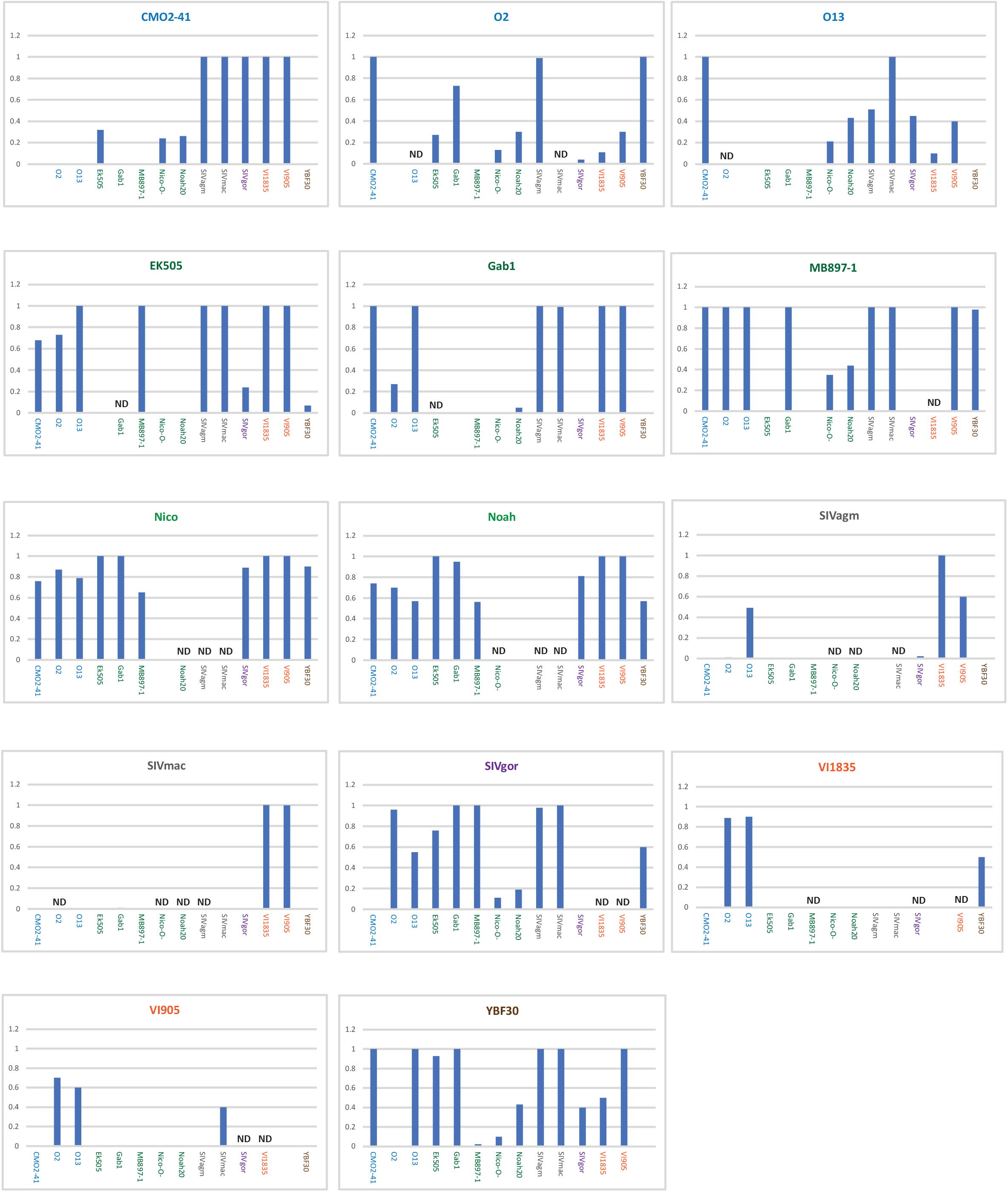
Fitness of primate lentiviruses in human PBMC. Primate lentivirus were added to *Hs*PBMCs for a full pairwise competition. Production of the “titled” virus in each chart is presented as fraction of the total when competed against the virus on the X axis. Relative virus production was measured by PCR and deep sequencing analyses using the SeekDeep pipeline. All competitions were performed in duplicate with a 10% observed variance in fitness.

**Supplementary Figure 3c:**
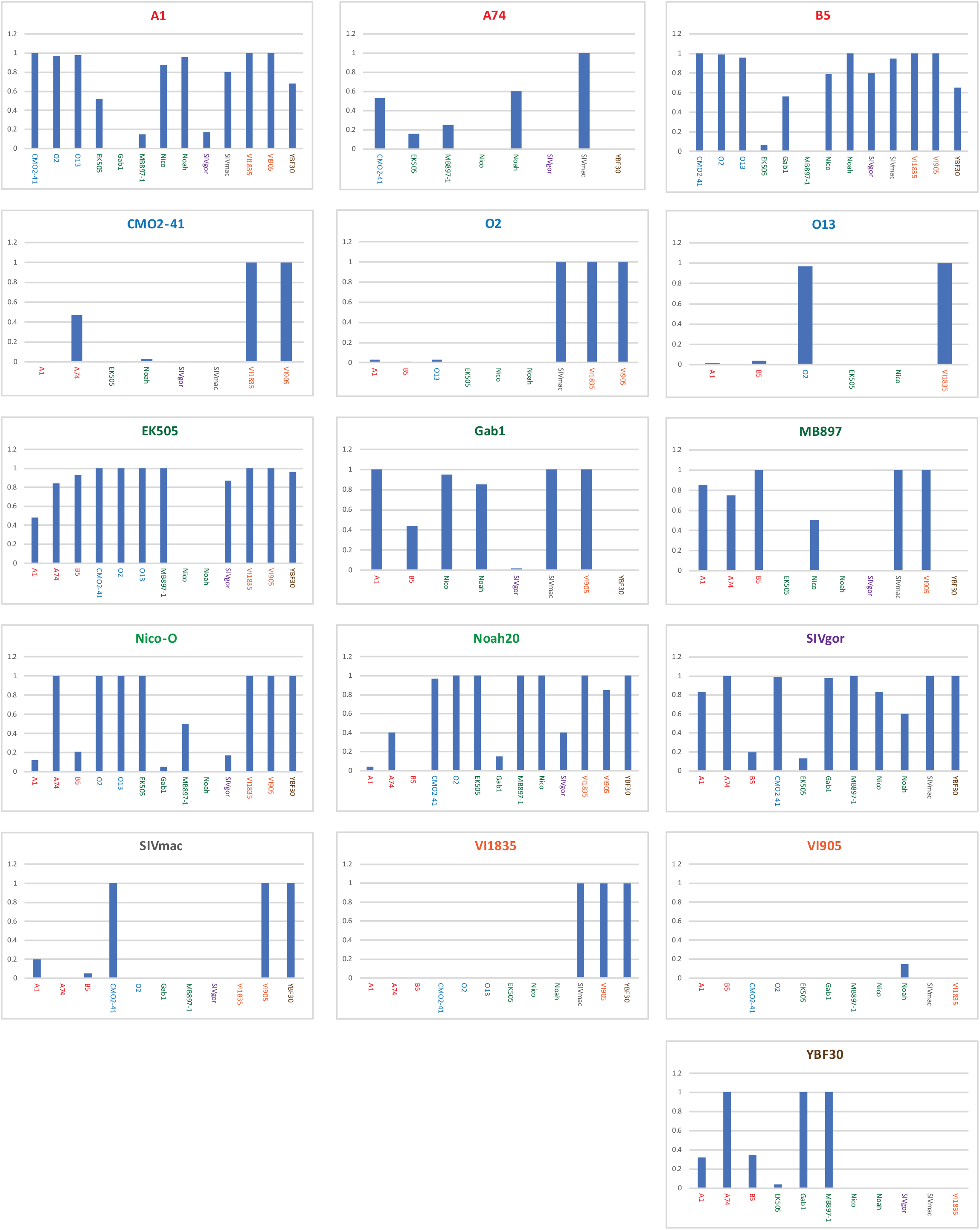
Fitness of primate lentiviruses in chimp PBMC. Primate lentivirus were added to *Ptv*PBMCs for a partial pairwise competition. Production of the “titled” virus in each chart is presented as fraction of the total when competed against the virus on the X axis. Relative virus production was measured from PCR and deep sequencing analyses using the SeekDeep pipeline. All competitions were performed in duplicate with a 10% observed variance in fitness.

**Supplementary Figure 4.**
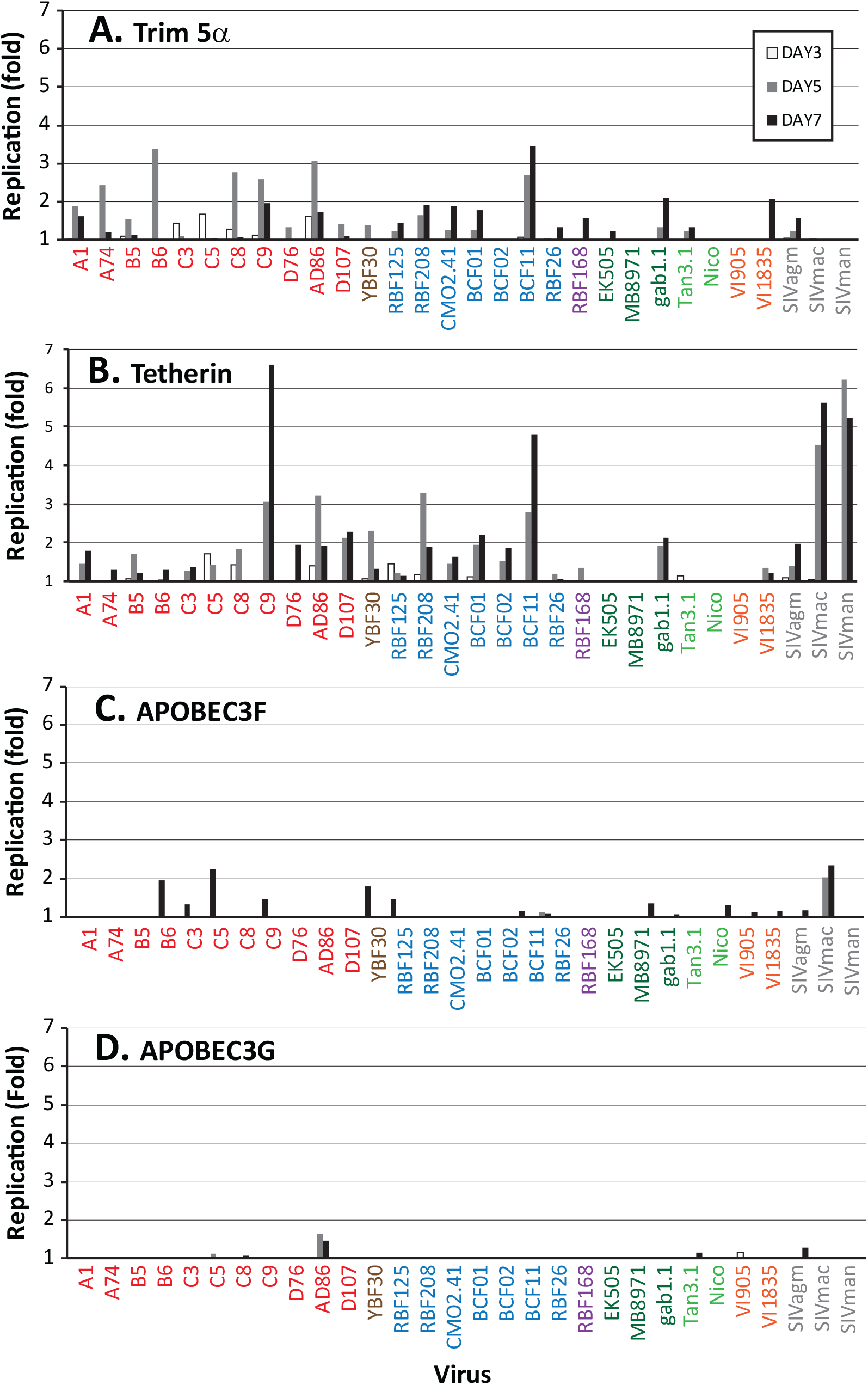
Primate lentivirus replication in human CD4+ U87 cells expressing HsCCR5 and pre-treated with siRNA to human restiriction factors Trim 5a (A), Tetherin/BST2 (B), APOBEC3F (C), and APOBEC3G (D). Replication is plotted as the fold change in increased lentivirus replication on days 3, 5, and 7 in the presence of siRNA to specific human restriction factors.

**Supplementary Table 1.** Statistics on lentiviral pairwise competitions in human PBMCs using group M HIV-1 as the compartor.

| Dunnett's multiple comparisons test | Mean Diff. | 95% CI of diff. | Significant? | Summary |
| --- | --- | --- | --- | --- |
| Group M vs. ALL | 0.09368 | 0.006571 to 0.1808 | Yes | p < 0.05 |
| Group M vs. SIVcpz | -0.02116 | -0.1400 to 0.09765 | No | ns |
| Group M vs. GrpO | 0.1563 | 0.01406 to 0.2985 | Yes | p < 0.05 |
| Group M vs. SIVgor | -0.14 | -0.3705 to 0.09046 | No | ns |
| Group M vs. GrpN | -0.09222 | -0.3227 to 0.1382 | No | ns |
| Group M vs. HIV-2 | 0.3547 | 0.1818 to 0.5276 | Yes | p < 0.0001 |
| Group M vs. SIVmac/smm | 0.2463 | 0.06867 to 0.4238 | Yes | p < 0.01 |

| Test details | Mean 1 | Mean 2 | Mean Diff. | SE of diff. | n1 | n2 | q | DF |
| --- | --- | --- | --- | --- | --- | --- | --- | --- |
| Group M vs. ALL | 0.5 | 0.4063 | 0.09368 | 0.03317 | 117 | 117 | 2.824 | 347 |
| Group M vs. SIVcpz | 0.5 | 0.5212 | -0.02116 | 0.04524 | 117 | 43 | 0.4678 | 347 |
| Group M vs. GrpO | 0.5 | 0.3437 | 0.1563 | 0.05416 | 117 | 27 | 2.886 | 347 |
| Group M vs. SIVgor | 0.5 | 0.64 | -0.14 | 0.08775 | 117 | 9 | 1.595 | 347 |
| Group M vs. GrpN | 0.5 | 0.5922 | -0.09222 | 0.08775 | 117 | 9 | 1.051 | 347 |
| Group M vs. HIV-2 | 0.5 | 0.1453 | 0.3547 | 0.06584 | 117 | 17 | 5.387 | 347 |
| Group M vs. SIVmac/smm | 0.5 | 0.2538 | 0.2463 | 0.06762 | 117 | 16 | 3.642 | 347 |

**Supplementary Table 2.** Statistics on lentiviral pairwise competitions in human PBMCs using SIVcpz as the compartor.

| Dunnett's multiple comparisons test | Mean Diff. | 95% CI of diff. | Significant? | Summary |
| --- | --- | --- | --- | --- |
| SIVcpz vs. ALL | 0.07132 | -0.03285 to 0.1755 | No | ns |
| SIVcpz vs. Grp M | 0.02393 | -0.1117 to 0.1596 | No | ns |
| SIVcpz vs. Grp O | 0.007857 | -0.2073 to 0.2230 | No | ns |
| SIVcpz vs. SIVgor | -0.112 | -0.4594 to 0.2354 | No | ns |
| SIVcpz vs. Grp N | 0.006 | -0.3414 to 0.3534 | No | ns |
| SIVcpz vs. HIV-2 | 0.5 | 0.2369 to 0.7631 | Yes | p < 0.0001 |
| SIVcpz vs. SIVmac/smm | 0.4833 | 0.1649 to 0.8017 | Yes | p < 0.001 |

| Test details | Mean 1 | Mean 2 | Mean Diff. | SE of diff. | n1 | n2 | q | DF |
| --- | --- | --- | --- | --- | --- | --- | --- | --- |
| SIVcpz vs. ALL | 0.5 | 0.4287 | 0.07132 | 0.03962 | 117 | 98 | 1.8 | 289 |
| SIVcpz vs. Grp M | 0.5 | 0.4761 | 0.02393 | 0.0516 | 117 | 43 | 0.4638 | 289 |
| SIVcpz vs. Grp O | 0.5 | 0.4921 | 0.007857 | 0.08183 | 117 | 14 | 0.09602 | 289 |
| SIVcpz vs. SIVgor | 0.5 | 0.612 | -0.112 | 0.1321 | 117 | 5 | 0.8476 | 289 |
| SIVcpz vs. Grp N | 0.5 | 0.494 | 0.006 | 0.1321 | 117 | 5 | 0.04541 | 289 |
| SIVcpz vs. HIV-2 | 0.5 | 0 | 0.5 | 0.1001 | 117 | 9 | 4.995 | 289 |
| SIVcpz vs. SIVmac/smm | 0.5 | 0.01667 | 0.4833 | 0.1211 | 117 | 6 | 3.991 | 289 |

**Supplementary Table 3.** Statistics on lentiviral pairwise competitions in chimp PBMCs using HIV-1 M as the compartor.

| Dunnett's multiple comparisons test | Mean Diff. | 95% CI of diff. | Significant? | Summary |
| --- | --- | --- | --- | --- |
| Group M vs. ALL | 0.1139 | 0.008974 to 0.2189 | Yes | p < 0.05 |
| Group M vs. SIVcpz | -0.08714 | -0.2377 to 0.06346 | No | ns |
| Group M vs. Grp O | 0.405 | 0.1821 to 0.6279 | Yes | p < 0.0001 |
| Group M vs. SIVgor | -0.1767 | -0.4880 to 0.1347 | No | ns |
| Group M vs. Grp N | 0.2433 | -0.06804 to 0.5547 | No | ns |
| Group M vs. HIV-2 | 0.5 | 0.3054 to 0.6946 | Yes | p < 0.0001 |
| Group M vs. SIVmac/man | 0.4167 | 0.1053 to 0.7280 | Yes | p < 0.01 |

| Test details | Mean 1 | Mean 2 | Mean Diff. | SE of diff. | n1 | n2 | q | DF |
| --- | --- | --- | --- | --- | --- | --- | --- | --- |
| Group M vs. ALL | 0.5 | 0.3861 | 0.1139 | 0.03978 | 117 | 33 | 2.864 | 179 |
| Group M vs. SIVcpz | 0.5 | 0.5871 | -0.08714 | 0.05707 | 117 | 14 | 1.527 | 179 |
| Group M vs. Grp O | 0.5 | 0.095 | 0.405 | 0.08448 | 117 | 6 | 4.794 | 179 |
| Group M vs. SIVgor | 0.5 | 0.6767 | -0.1767 | 0.118 | 117 | 3 | 1.497 | 179 |
| Group M vs. Grp N | 0.5 | 0.2567 | 0.2433 | 0.118 | 117 | 3 | 2.062 | 179 |
| Group M vs. HIV-2 | 0.5 | 0 | 0.5 | 0.07375 | 117 | 8 | 6.779 | 179 |
| Group M vs. SIVmac/man | 0.5 | 0.08333 | 0.4167 | 0.118 | 117 | 3 | 3.531 | 179 |

**Supplementary Table 4.**
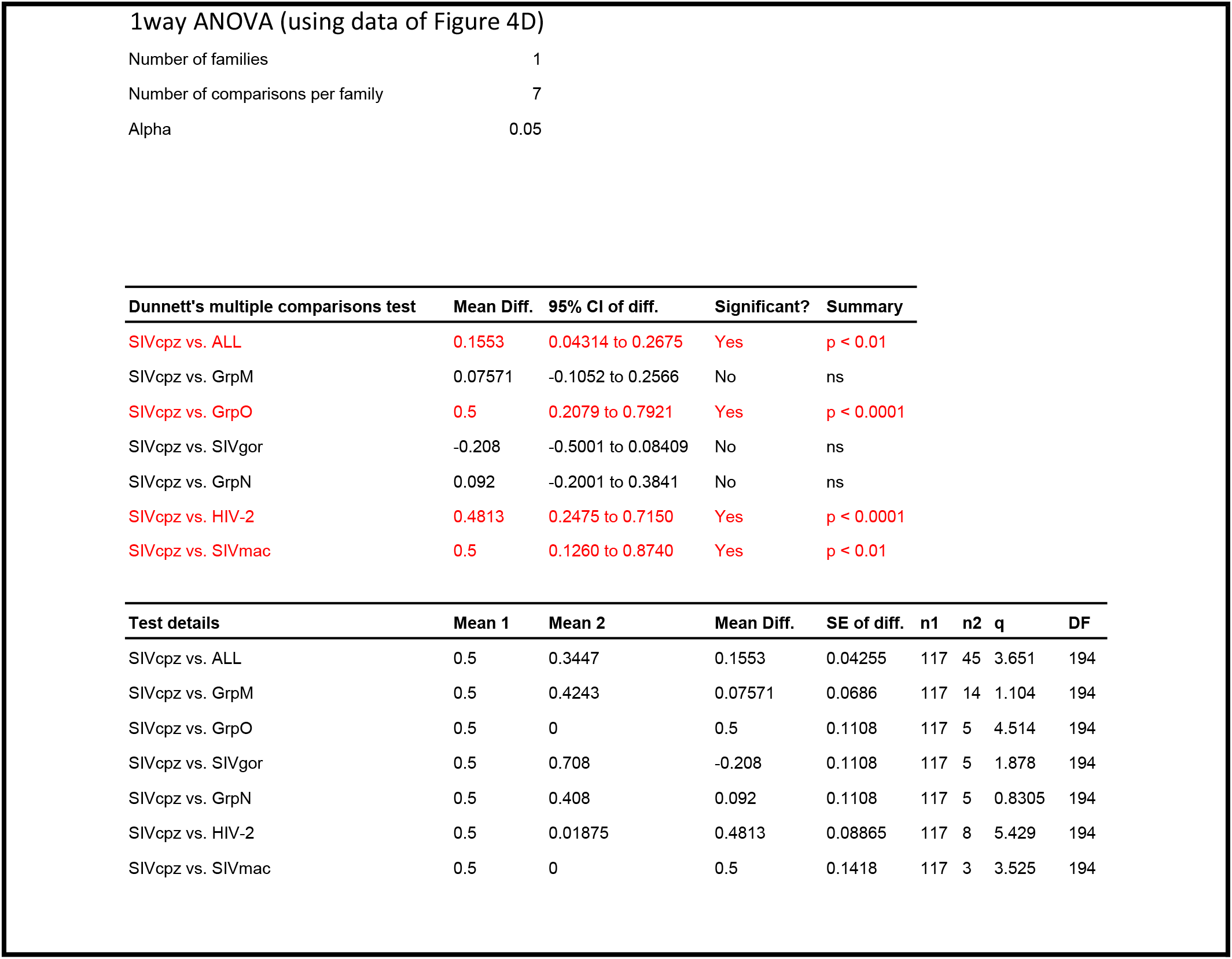
Statistics on lentiviral pairwise competitions in chimp PBMCs using SIVcpz as the compartor.

**Supplementary Table 5.** Statistics on replicative fitness of specific group/type against intergroup/type in human PBMCs.

| Tukey's multiple comparisons test | Mean Diff. | 95% CI of diff. | Significant? | Summary |
| --- | --- | --- | --- | --- |
| group M vs. group O | 0.1247 | -0.05142 to 0.3009 | No | ns |
| group M vs. group N | 0.02411 | -0.2330 to 0.2812 | No | ns |
| group M vs. SIVcpz | 0.01007 | -0.1475 to 0.1676 | No | ns |
| group M vs. SIVcpz_ptt | 0.007242 | -0.1767 to 0.1912 | No | ns |
| group M vs. SIVcpz_pts | 0.01467 | -0.2058 to 0.2352 | No | ns |
| group M vs. SIVgor | -0.03389 | -0.3017 to 0.2339 | No | ns |
| group M vs. HIV-2 | 0.4066 | 0.1982 to 0.6150 | Yes | p < 0.0001 |
| group M vs. SIVagm/smm | 0.383 | 0.1676 to 0.5983 | Yes | p < 0.0001 |
| group O vs. group N | -0.1006 | -0.3777 to 0.1765 | No | ns |
| group O vs. SIVcpz | -0.1147 | -0.3031 to 0.07378 | No | ns |
| group O vs. SIVcpz_ptt | -0.1175 | -0.3285 to 0.09352 | No | ns |
| group O vs. SIVcpz_pts | -0.1101 | -0.3536 to 0.1335 | No | ns |
| group O vs. SIVgor | -0.1586 | -0.4457 to 0.1284 | No | ns |
| group O vs. HIV-2 | 0.2819 | 0.04924 to 0.5145 | Yes | p < 0.01 |
| group O vs. SIVagm/smm | 0.2582 | 0.01938 to 0.4971 | Yes | p < 0.05 |
| group N vs. SIVcpz | -0.01404 | -0.2797 to 0.2517 | No | ns |
| group N vs. SIVcpz_ptt | -0.01687 | -0.2990 to 0.2653 | No | ns |
| group N vs. SIVcpz_pts | -0.009439 | -0.3167 to 0.2978 | No | ns |
| group N vs. SIVgor | -0.05801 | -0.4008 to 0.2847 | No | ns |
| group N vs. HIV-2 | 0.3825 | 0.08383 to 0.6812 | Yes | p < 0.01 |
| group N vs. SIVagm/smm | 0.3589 | 0.05532 to 0.6624 | Yes | p < 0.01 |
| SIVcpz vs. SIVcpz_ptt | -0.002831 | -0.1986 to 0.1929 | No | ns |
| SIVcpz vs. SIVcpz_pts | 0.0046 | -0.2259 to 0.2351 | No | ns |
| SIVcpz vs. SIVgor | -0.04397 | -0.3200 to 0.2321 | No | ns |
| SIVcpz vs. HIV-2 | 0.3965 | 0.1776 to 0.6154 | Yes | p < 0.0001 |
| SIVcpz vs. SIVagm/smm | 0.3729 | 0.1474 to 0.5984 | Yes | p < 0.0001 |
| SIVcpz_ptt vs. SIVcpz_pts | 0.007431 | -0.2418 to 0.2567 | No | ns |
| SIVcpz_ptt vs. SIVgor | -0.04114 | -0.3330 to 0.2508 | No | ns |
| SIVcpz_ptt vs. HIV-2 | 0.3994 | 0.1608 to 0.6380 | Yes | p < 0.0001 |
| SIVcpz_ptt vs. SIVagm/smm | 0.3757 | 0.1311 to 0.6204 | Yes | p < 0.0001 |
| SIVcpz_pts vs. SIVgor | -0.04857 | -0.3648 to 0.2677 | No | ns |
| SIVcpz_pts vs. HIV-2 | 0.3919 | 0.1241 to 0.6597 | Yes | p < 0.0001 |
| SIVcpz_pts vs. SIVagm/smm | 0.3683 | 0.09507 to 0.6415 | Yes | p < 0.01 |
| SIVgor vs. HIV-2 | 0.4405 | 0.1326 to 0.7484 | Yes | p < 0.001 |
| SIVgor vs. SIVagm/smm | 0.4169 | 0.1042 to 0.7295 | Yes | p < 0.01 |
| HIV-2 vs. SIVagm/smm | -0.02363 | -0.2872 to 0.2399 | No | ns |

| Test details | Mean 1 | Mean 2 | Mean Diff. | SE of diff. | n1 | n2 | q | DF |
| --- | --- | --- | --- | --- | --- | --- | --- | --- |
| group M vs. group O | 0.602 | 0.4773 | 0.1247 | 0.05652 | 121 | 59 | 3.121 | 452 |
| group M vs. group N | 0.602 | 0.5779 | 0.02411 | 0.08249 | 121 | 22 | 0.4134 | 452 |
| group M vs. SIVcpz | 0.602 | 0.592 | 0.01007 | 0.05055 | 121 | 84 | 0.2818 | 452 |
| group M vs. SIVcpz_ptt | 0.602 | 0.5948 | 0.007242 | 0.05902 | 121 | 52 | 0.1735 | 452 |
| group M vs. SIVcpz_pts | 0.602 | 0.5874 | 0.01467 | 0.07075 | 121 | 32 | 0.2933 | 452 |
| group M vs. SIVgor | 0.602 | 0.6359 | -0.03389 | 0.08591 | 121 | 20 | 0.5579 | 452 |
| group M vs. HIV-2 | 0.602 | 0.1954 | 0.4066 | 0.06686 | 121 | 37 | 8.6 | 452 |
| group M vs. SIVagm/smm | 0.602 | 0.2191 | 0.383 | 0.06909 | 121 | 34 | 7.84 | 452 |
| group O vs. group N | 0.4773 | 0.5779 | -0.1006 | 0.08891 | 59 | 22 | 1.6 | 452 |
| group O vs. SIVcpz | 0.4773 | 0.592 | -0.1147 | 0.06046 | 59 | 84 | 2.682 | 452 |
| group O vs. SIVcpz_ptt | 0.4773 | 0.5948 | -0.1175 | 0.0677 | 59 | 52 | 2.454 | 452 |
| group O vs. SIVcpz_pts | 0.4773 | 0.5874 | -0.1101 | 0.07814 | 59 | 32 | 1.992 | 452 |
| group O vs. SIVgor | 0.4773 | 0.6359 | -0.1586 | 0.09209 | 59 | 20 | 2.436 | 452 |
| group O vs. HIV-2 | 0.4773 | 0.1954 | 0.2819 | 0.07464 | 59 | 37 | 5.341 | 452 |
| group O vs. SIVagm/smm | 0.4773 | 0.2191 | 0.2582 | 0.07664 | 59 | 34 | 4.766 | 452 |
| group N vs. SIVcpz | 0.5779 | 0.592 | -0.01404 | 0.08524 | 22 | 84 | 0.2329 | 452 |
| group N vs. SIVcpz_ptt | 0.5779 | 0.5948 | -0.01687 | 0.09052 | 22 | 52 | 0.2636 | 452 |
| group N vs. SIVcpz_pts | 0.5779 | 0.5874 | -0.009439 | 0.09857 | 22 | 32 | 0.1354 | 452 |
| group N vs. SIVgor | 0.5779 | 0.6359 | -0.05801 | 0.11 | 22 | 20 | 0.746 | 452 |
| group N vs. HIV-2 | 0.5779 | 0.1954 | 0.3825 | 0.09582 | 22 | 37 | 5.645 | 452 |
| group N vs. SIVagm/smm | 0.5779 | 0.2191 | 0.3589 | 0.09739 | 22 | 34 | 5.211 | 452 |
| SIVcpz vs. SIVcpz_ptt | 0.592 | 0.5948 | -0.002831 | 0.0628 | 84 | 52 | 0.06375 | 452 |
| SIVcpz vs. SIVcpz_pts | 0.592 | 0.5874 | 0.0046 | 0.07394 | 84 | 32 | 0.08799 | 452 |
| SIVcpz vs. SIVgor | 0.592 | 0.6359 | -0.04397 | 0.08856 | 84 | 20 | 0.7021 | 452 |
| SIVcpz vs. HIV-2 | 0.592 | 0.1954 | 0.3965 | 0.07023 | 84 | 37 | 7.985 | 452 |
| SIVcpz vs. SIVagm/smm | 0.592 | 0.2191 | 0.3729 | 0.07235 | 84 | 34 | 7.289 | 452 |
| SIVcpz_ptt vs. SIVcpz_pts | 0.5948 | 0.5874 | 0.007431 | 0.07997 | 52 | 32 | 0.1314 | 452 |
| SIVcpz_ptt vs. SIVgor | 0.5948 | 0.6359 | -0.04114 | 0.09365 | 52 | 20 | 0.6212 | 452 |
| SIVcpz_ptt vs. HIV-2 | 0.5948 | 0.1954 | 0.3994 | 0.07655 | 52 | 37 | 7.378 | 452 |
| SIVcpz_ptt vs. SIVagm/smm | 0.5948 | 0.2191 | 0.3757 | 0.0785 | 52 | 34 | 6.769 | 452 |
| SIVcpz_pts vs. SIVgor | 0.5874 | 0.6359 | -0.04857 | 0.1015 | 32 | 20 | 0.677 | 452 |
| SIVcpz_pts vs. HIV-2 | 0.5874 | 0.1954 | 0.3919 | 0.08592 | 32 | 37 | 6.451 | 452 |
| SIVcpz_pts vs. SIVagm/smm | 0.5874 | 0.2191 | 0.3683 | 0.08766 | 32 | 34 | 5.942 | 452 |
| SIVgor vs. HIV-2 | 0.6359 | 0.1954 | 0.4405 | 0.09878 | 20 | 37 | 6.307 | 452 |
| SIVgor vs. SIVagm/smm | 0.6359 | 0.2191 | 0.4169 | 0.1003 | 20 | 34 | 5.878 | 452 |
| HIV-2 vs. SIVagm/smm | 0.1954 | 0.2191 | -0.02363 | 0.08456 | 37 | 34 | 0.3953 | 452 |

**Supplementary Table 6.**
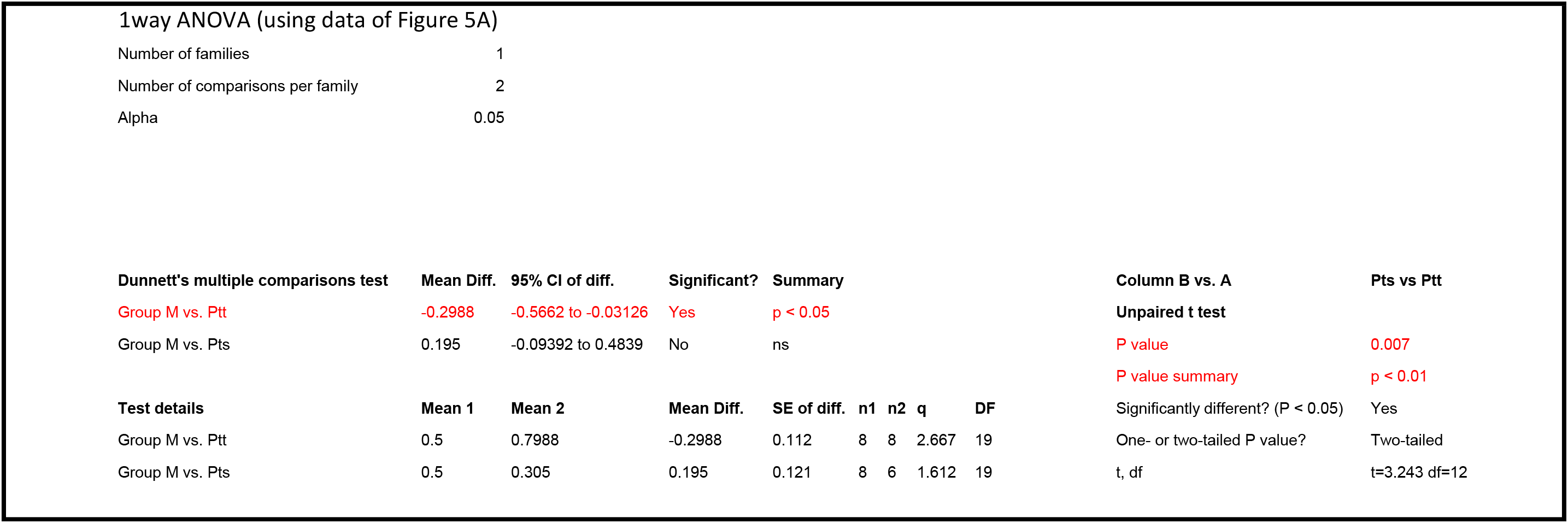
Statistics on SIVcpz_Ptt and SIVcpz_Pts pairwise competitions in human PBMCs using HIV-1 M as the compartor.

1way ANOVA (using data of Figure 5A)
|  |  |
| --- | --- |
| Number of families | 1 |
| Number of comparisons per family | 2 |
| Alpha | 0.05 |

| Dunnett's multiple comparisons test | Mean Diff. | 95% CI of diff. | Significant? | Summary | Column B vs. A | Pts vs Ptt |  |  |  |  |
| --- | --- | --- | --- | --- | --- | --- | --- | --- | --- | --- |
| Group M vs. Ptt | -0.2988 | -0.5662 to -0.03126 | Yes | p < 0.05 | Unpaired t test |  |  |  |  |  |
| Group M vs. Pts | 0.195 | -0.09392 to 0.4839 | No | ns | P value | 0.007 |  |  |  |  |
|  |  |  |  |  | P value summary | p < 0.01 |  |  |  |  |
| Test details | Mean 1 | Mean 2 | Mean Diff. | SE of diff. | n1 | n2 | q | DF | Significantly different? (P < 0.05) | Yes |
| Group M vs. Ptt | 0.5 | 0.7988 | -0.2988 | 0.112 | 8 | 8 | 2.667 | 19 | One- or two-tailed P value? | Two-tailed |
| Group M vs. Pts | 0.5 | 0.305 | 0.195 | 0.121 | 8 | 6 | 1.612 | 19 | t, df | t=3.243 df=12 |

**Supplementary Table 7.**
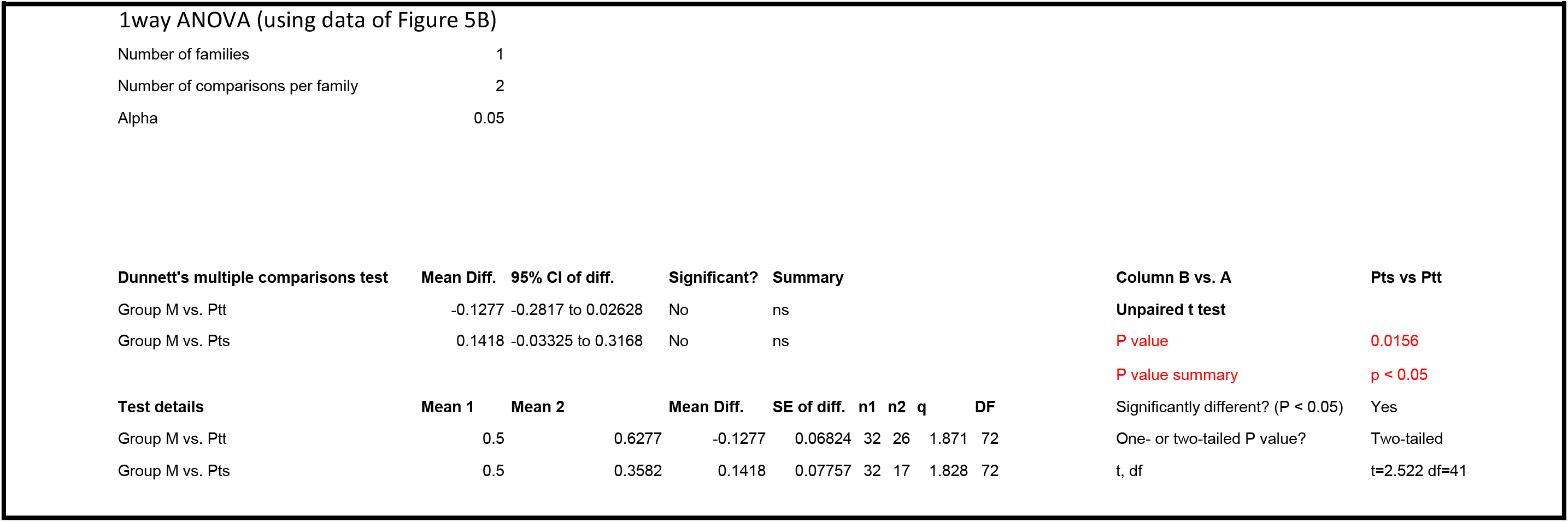
Statistics on SIVcpz_Ptt and SIVcpz_Pts pairwise competitions in chimp PBMCs using HIV-1 M as the compartor.

| Dunnett's multiple comparisons test | Mean Diff. | 95% CI of diff. | Significant? | Summary | Column B vs. A | Pts vs Ptt |  |  |  |  |
| --- | --- | --- | --- | --- | --- | --- | --- | --- | --- | --- |
| Group M vs. Ptt | -0.1277 | -0.2817 to 0.02628 | No | ns | Unpaired t test |  |  |  |  |  |
| Group M vs. Pts | 0.1418 | -0.03325 to 0.3168 | No | ns | P value | 0.0156 |  |  |  |  |
|  |  |  |  |  | P value summary | p < 0.05 |  |  |  |  |
| Test details | Mean 1 | Mean 2 | Mean Diff. | SE of diff. | n1 | n2 | q | DF | Significantly different? (P < 0.05) | Yes |
| Group M vs. Ptt | 0.5 | 0.6277 | -0.1277 | 0.06824 | 32 | 26 | 1.871 | 72 | One- or two-tailed P value? | Two-tailed |
| Group M vs. Pts | 0.5 | 0.3582 | 0.1418 | 0.07757 | 32 | 17 | 1.828 | 72 | t, df | t=2.522 df=41 |

